# Deleting *in vivo* β-catenin degradation domain in mouse hepatocytes drives hepatocellular carcinoma or hepatoblastoma-like tumors

**DOI:** 10.1101/2021.07.04.450836

**Authors:** Robin Loesch, Stefano Caruso, Valérie Paradis, Cecile Godard, Angélique Gougelet, Simon Picard, Christine Perret, Makoto Mark Taketo, Jessica Zucman-Rossi, Sabine Colnot

## Abstract

**Background and aims:** One-third of hepatocellular carcinomas (HCCs) have mutations that activate the β-catenin pathway with mostly *CTNNB1* mutations. Mouse models using *Adenomatous polyposis coli* (*Apc*) loss-of-functions (LOF) are widely used to mimic β-catenin-dependent tumorigenesis. Considering the low prevalence of *APC* mutations in human HCCs we aimed to generate hepatic tumors through *CTNNB1* exon 3 deletion (β*cat*^Δex3^) and to compare them to hepatic tumors with *Apc* LOF engineered through a frameshift in exon 15 (*Apc^fs-ex15^*).

**Methods:** We used hepatic-specific and inducible Cre-lox mouse models as well as a hepatic-specific *in vivo* CRISPR/Cas9 approach using AAV vectors, to generate *Apc^fs-ex15^* and βcat^Δex3^ hepatic tumors harboring activation of the β-catenin pathway. Tumors generated by the Cre-lox models were analyzed phenotypically using immunohistochemistry and were selected for transcriptomic analysis using RNA-sequencing. Mouse RNAseq data were compared to human RNAseq data (normal tissues (8), HCCs (48) and hepatoblastomas (9)) in an integrative analysis. Tumors generated via CRISPR were analyzed using DNA sequencing and immunohistochemistry.

**Results:** Mice with βcat^Δex3^ alteration in hepatocytes developed liver tumors. Generated tumors were indistinguishable from those arising in *Apc^fs-ex15^* mice. Both *Apc^fs-ex15^* and βcat^Δex3^ mouse models induced two phenotypically distinct tumors (differentiated or undifferentiated). Integrative analysis of human and mouse tumors showed that mouse differentiated tumors are close to human well differentiated *CTNNB1*-mutated tumors, while undifferentiated ones are closer to human mesenchymal hepatoblastomas, and are activated for YAP signaling.

**Conclusion:** *Apc^fs-ex15^* and βcat^Δex3^ mouse models similarly induce tumors transcriptionally close to either well differentiated β-Catenin activated human HCCs or mesenchymal hepatoblastomas.

## Introduction

Hepatocellular carcinoma (HCC), the most frequent liver primary cancer, is the fourth highest cause of cancer-related death^1^. It mainly arises on a diseased liver, associated with hepatitis B (HBV) and C (HCV) viral infections, alcohol abuse or metabolic syndrome. HCC development relies on a multistep process involving genetic alterations. Prevailing mutational hotspots highlight telomerase activation, TP53 or β-catenin signaling, and chromatin remodeling process, as being the core pathways supporting HCC pathogenesis. The oncogenic property of dysregulated Wnt/β-Catenin pathway in HCC was first suggested after the discovery of activating somatic point mutations in *CTNNB1* gene encoding β-Catenin both in human and mouse HCC^2^. Nowadays, *CTNNB1* mutations in human HCC are well described and occur mainly in its exon 3, disrupting the degradation domain of β-catenin. Mutations lead to β-catenin stabilization, to its translocation in the nucleus where it participates to a transcriptional complex with the LEF/TCF nuclear factors, regulating gene expression. Some target genes of nuclear hepatic β-catenin such as *GLUL* encoding Glutamine Synthetase (GS) and *AXIN2*, are markers of *CTNNB1*-mutated human HCCs^3, 4^. Even though the majority of the mutations that activate the β-Catenin pathway are found in *CTNNB1* itself (37% of HCCs), mutations in *Adenomatous Polyposis Coli* (*APC)* and *AXIN1*, which are involved in β-catenin degradation complex, are also found in 1-2% and 11% of HCCs respectively^5^. A recent study showed that *AXIN1* deficiency in human and mouse induces HCC without β-Catenin activation^6^. Conversely, we previously described the inducible and liver-specific loss of function of *Apc*, obtained through a Cre-loxP mediated excision of its exon 14, generating a frameshift in exon 15, thereafter called the *Apc*^fs-ex15^ mouse model: *Apc* loss in single hepatocytes led to β-catenin signaling and liver tumorigenesis^7^. Since *APC* mutations are found in only 1% of human HCCs, we could not exclude that β-catenin dependent tumorigenesis in the *Apc*^fs-ex15^ mouse does not phenocopy that linked to *CTNNB1* gene alterations. It could be due to multiple alternative functions exerted by APC, including its role in chromosomal instability found in colorectal cancers^8^.

About 10% of *CTNNB1* mutations found in human HCCs or hepatocellular adenomas (HCA) are large deletions in exon 3 leading to in-frame deletion of sequences encoding the residues whose phosphorylation is needed for β-Catenin degradation (serine 33-37-45 and threonine 41)^4^. Hepatoblastoma is a pediatric liver primary cancer in which *CTNNB1* is also a key driver gene through in-frame mutation (50-90%) mainly consisting in deletions surrounding the exon 3^9^. In 2002, Harada *et al* generated a mouse model generating a Cre-loxP-mediated hepatic loss of the exon 3 of *Ctnnb1* (β*cat^Δex3^*), in which no liver tumorigenesis occurred 6 months after adenovirus-CRE (AdCre) injection^10^. Here we used an alternate tamoxifen-inducible approach for Cre recombination to test liver tumorigenesis in β*cat^Δex3^* mice, similar to a protocol efficient for inducing liver tumorigenesis in *Apc*^fs-ex15^ mice^3, 11–13^. We also edited *Apc* or *Ctnnb1* in hepatocytes by performing *in vivo* an AAV-mediated delivery of CRISPR/Cas9 machinery^14^.

We showed that both Cre-loxP- and CRISPR-Cas9-mediated β*cat^Δex3^* alteration in hepatocytes induce β-catenin-dependent liver tumorigenesis similarly to the *Apc*^fs-ex15^ mouse model. Regardless of the Wnt partner targeted and of the gene editing strategy, the mice generated two phenotypically distinct types of tumors that we defined as well-differentiated (DIFF) or poorly-differentiated (UNDIFF) tumors. Integrative analysis of RNAseq data from mouse tumors with human HCCs or hepatoblastomas revealed that DIFF tumors in mouse are close to well differentiated *CTNNB1*-mutated human HCCs, while UNDIFF mouse tumors are close to human mesenchymal hepatoblastomas.

## Material and methods

### In vivo CRISPR design and editing analysis

All the guides used were designed using CRISPR RGEN online tool (Cas-Designer http://www.rgenome.net/cas-designer/). Twenty-one nucleotides long guides were selected according to their GC content (20% to 80%), low potential off-target sites and position in previously tested PCR for easy and efficient indels analysis via T7 nuclease assay or SANGER sequencing followed by online analysis by TIDE (Tracking of Indels by DEcomposition; https://tide.deskgen.com/). Guides were cloned in single AAV-Cas9 plasmids containing both sgRNA and Cas9 from *Staphylococcus aureus,* driven by either a CMV promoter (pX601, addgene #61591, used for *in vitro* studies), or a TBG promoter (pX602, addgene #61593, Figure 2A)^14^. Plasmids were then transfected independently into mouse hepatoma Hepa1-6 cells for *Apc* and *Rosa26* guides, or mouse AML12 cell line for the βcat^Δex3^ strategy. For each locus, three guides were tested *in vitro* to select the most efficient ones (**Table S1**). Two days after transfection, cell DNA was extracted and amplified by PCR for the targeted loci.

### Cell culture and transfection

Hepa1-6 cells were maintained in DMEM medium supplemented with 10% fetal calf serum and 1% penicillin-streptomycin. AML12 cells were maintained in DMEM/F12 medium (1:1) supplemented with 10% fetal calf serum and 1% penicillin-streptomycin. Hepa1-6 and AML12 cells were transfected in six well plates using 10μl lipofectamine 2000 (Life Technologies). Due to a mutation deleting a part of *Ctnnb1* exon 3 in Hepa1-6 (**Figure S5C**), AML12 cells were used to test the guide RNAs targeting intron 2 and 3.

### Editing analysis

The PCR product of the targeted loci were sequenced by Sanger for assessment of total DNA insertions and deletions (indels), using TIDE or analyzed by a T7 nuclease assay (Surveyor^®^ Mutation Detection Kit – IDT 706020). Surveyor assays were performed for *Apc* and *Rosa26* initial experiments only, as it appeared more quantitative and reproducible to use Tide. For Sanger sequencing, the DNA was extracted on gel after PCR. Guides and PCR primers are in **Table S1**. To measure *in vivo* the efficiency of *Ctnnb1* exon3 editing, specific PCRs were performed from βcat^Δex3^ livers and tumors: Taqman assays (Thermo Fisher Scientific) amplified either WT *Ctnnb1* (Mm00483025), or WT and exon 3-deleted *Ctnnb1* (Mm01350394).

### Production of AAV-CRISPR/Cas9 viral particles

pX602 plasmids were amplified in Stbl3 bacterial strain then purified using Qiagen Plasmid Maxi Kit (ID:12162) and sent to the Center of viral vector production at the Health Research Institute in Nantes for recombinant AAV8 production (CPV INSERM UMR1089 Université de Nantes).

### Animal injection and processing

All animal procedures were approved by the ethical committee of Université de Paris according to the French government regulation (APAFIS #16420). The mice were maintained at the animal facility with standard diet and housing. Retro-orbital injections, ultrasonography and liver biopsies were done under general anesthesia with isoflurane inhalation. The *Apc*^fs-ex15^ mouse model was generated from compound *Apc*^flox/flox^/TTR-Cre^Tam^ injected with 0.75 mg Tamoxifen via intraperitoneal injection after dilution in oil^12, 13^. βcat^exon3-flox^ mice were intercrossed with TTR-Cre^Tam^ mice to generate a tamoxifen inducible and hepatocyte-specific model for *Ctnnb1* exon 3 deletion (βcat^Δex3^)10,^15^. Two month-old βcat^exon3-flox^/TTR-Cre^Tam^ male mice were also injected with Tamoxifen. Alternatively, they were given a tamoxifen diet (M-Z, low phytoestrogen + 400 mg/kg TAM citrate, SSNIFF, Soest, Germany) for two days after an overnight fasting. For the CRISPR-Cas9 strategy, retro-orbital injections of AAVs diluted in 200μl of physiological serum were done in 2-month-old C57Bl6/N male mice. Liver biopsies of the ligated extremity of the left median lobe (0.1cm^3^) were performed after minimal laparotomy. Excepted for the Cre-loxP βcat^Δex3^ mice, *Apc*^fs-ex15^ and βcat^Δex3^ mice were followed by ultrasonography every month until tumor detection, thereafter ultrasound imaging was continued every 2 weeks, as described^12^. Mice were sacrificed when tumor mass exceeded 2cm^3^ par mouse. The lethality rates indicate either a sudden death, or a sacrifice when ethical endpoints were reached.

### Immunohistochemistry (IHC) and staining

Hematoxylin-Eosin stainings and IHC were done on Formalin-fixed paraffin-embedded (FFPE) liver tissue sections as previously described2. The antibodies used are in **Table S2**.

### RNA and DNA extraction

Total RNAs were extracted from tumor or non-tumor tissue with Trizol reagent (ThermoFisher Scientific – 15596018) as previously described^12^. DNA extractions were performed using classical phenol-chloroform protocol (Invitrogen Ultrapure Phenol:Chloroform:Isoamyl Alcohol (25:24:1, v/v)).

### Mouse RNA sequencing

Libraries were prepared by Integragen SA with NEBNext Ultra II Directional RNA Library Prep Kit for Illumina protocol according supplier recommendations. Briefly the key stages of this protocol are successively, the purification of PolyA containing mRNA molecules using poly-T oligo attached magnetic beads from 1μg total RNA (with the Magnetic mRNA Isolation Kit from NEB), a fragmentation using divalent cations under elevated temperature to obtain approximately 300bp pieces, double strand cDNA synthesis and finally Illumina adapters ligation and cDNA library amplification by PCR for sequencing. Sequencing was then carried out on paired-end 75 b of Illumina HiSeq4000. Base calling was performed by using the Real-Time Analysis software sequence pipeline (2.7.7) with default parameters. Quality of reads was assessed for each sample using FastQC (V.0.11.4; http://www.bioinformatics.babraham.ac.uk/projects/fastqc/). Mouse RNAseq analysis was performed using Galileo online software provided by Integragen. Data have been deposited in the European Nucleotide Archive (ENA) at EMBL-EBI under accession number PRJEB44400. (https://www.ebi.ac.uk/ena/browser/view/PRJEB44400)

### Human-Mouse integrative analysis

Statistical analysis and data visualization were performed using R software version 3.5.1 (R Foundation for Statistical Computing, Vienna, Austria. https://www.R-project.org) and Bioconductor version 3.4. A total of 80 samples from 2 different RNA-sequencing datasets was used for the comparative transcriptomic analysis, including 65 human and 15 mouse samples, respectively. The two datasets were initially processed separately. For each dataset, FPKM scores (number of fragments per kilobase of exon model and millions of mapped reads) were calculated by normalizing the count matrix for the library size and the coding length of each gene. FPKMs were, subsequently, log-transformed after adding a pseudocount of 1. First, we filtered out most of the genes by keeping only the 1000 genes with the highest variance in both datasets and, then, only common genes that are 1:1 orthologs in human and mouse were selected (n=324) using the list of mouse-human 1:1 orthologous genes from MGI (http://www.informatics.jax.org). Finally, we standardized gene expression separately to have mean 0 and standard deviation 1 per gene just before the two datasets were integrated. Unsupervised hierarchical clustering of the integrated data was performed using Pearson distance and Ward’s linkage method with ComplexHeatmap package in R. Principal component analysis (PCA) using the main two components was also performed. Pairwise correlations between mouse and human samples were determined using the expression profiles of the 324 mouse-human 1:1 orthologous genes and the Corrplot package in R. Hierarchical clustering of the obtained correlation coefficients was done using euclidean distance and average linkage method.

## Results

### βcat^Δex3^ CreloxP-engineered gene alteration in mouse induces liver cancer

We first asked whether βcat^Δex3^ mice develop liver tumors, as Apc^fs-ex15^ mice do. Lethality had been shown to occur when βcat^exon3-flox^ or Apcflox mice are injected with a high dose of Cre-Adenovirus (AdCre), as most of the hepatocytes are β-catenin-activated^7, 10^. We previously showed that a 2-fold lower dose of AdCre was sufficient to delete *Apc*, leading to β-catenin activation in single hepatocytes, and to tumorigenesis within 9 months^7^. A 10-fold lower dose of AdCre had been previously used for βcat^exon3-flox^ mice: this dose activated β-catenin in rare hepatocytes and was not tumorigenic 6 months after injection^10^. We therefore decided to implement an alternate protocol to activate β-catenin in βcat^exon3-flox^/TTR-Cre^Tam^ mouse livers, using Tamoxifen to activate Cre recombinase from TTR-Cre^Tam^ transgene^15^. We also performed a longer follow-up of tumors, until 12 months after Tamoxifen treatment.

When βcat^exon3-flox^/TTR-Cre^Tam^ mice were injected with 2 mg Tamoxifen, all the hepatocytes excepted the most periportal ones were β-catenin-activated (**Figure S1**). To activate β-catenin signaling in single hepatocytes and study tumorigenesis, βcat^exon3-flox^/TTR-Cre^Tam^ mice were either injected with 1 mg or 0.5 mg tamoxifen, or were given a tamoxifen-enriched diet for 2 days (**Figure 1**). High lethality still occurred in mice within one month after injection with 1 mg or 0.5 mg tamoxifen, with the long term survival of 23% and 58% of the mice respectively. Feeding mice with tamoxifen-enriched diet for 2 days increased mice viability to 90%.

**Figure 1:**
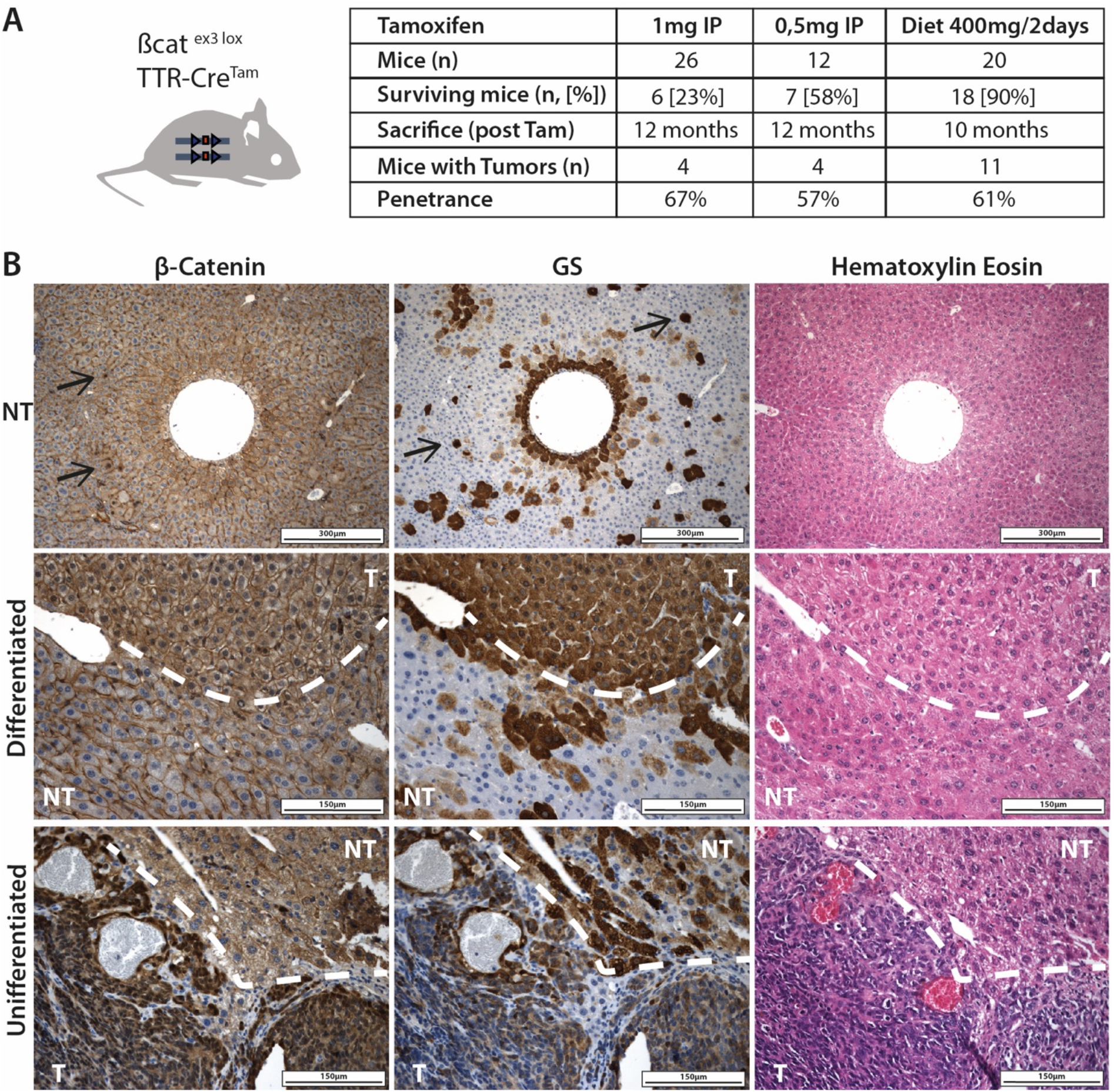
Emergence of differentiated and undifferentiated tumors after Tamoxifen-induced hepatic-specific loss of β-catenin-exon 3. **A:** Mice used in the study; **B:** Representative hematoxyline-Eosine stainings and IHC for β-catenin, glutamine synthetase (GS), on non-tumoral tissue (NT) and tumoral tissue. Black arrows: β-catenin activated hepatocytes.

Nineteen out of 31 surviving βcat^Δex3^ mice developed hepatic tumors during the 12 months following tamoxifen exposure. The status of the β-catenin signaling in tumors (T) and non-tumor (NT) tissue was assessed using β-catenin and glutamine synthetase (GS) staining (**Figure 1, Figure S6**). Cytoplasmic and/or nuclear localization of β-catenin as well as GS expression was found in few hepatocytes in NT liver. Like our previous observations with *Apc*^fs-ex15^ mouse model, we observed the development of two types of tumors, either well-differentiated (DIFF) or poorly-differentiated (UNDIFF)^7^. Well-differentiated tumors were characterized by hepatocyte-like tumor cells that maintain the β-catenin-induced expression of GS and present slight cytosolic or nuclear β-Catenin. Undifferentiated tumors were composed of small cells with basophilic nuclei. They strongly express nuclear β-Catenin but lose the expression of GS which is reminiscent of a loss of differentiation **(Figure 1, Figure S6)**. Among nine isolated tumors, 5 were classified as DIFF and 4 as UNDIFF (**Figure 3A**). The data clearly emphasize that a dominant stable mutant of β-catenin is sufficient to induce liver carcinogenesis in mouse.

### βcat^Δex3^ and Apc^fs-ex15^ mouse models can be generated through CRISPR/Cas9

In order to implement an easy-to-use model of βcat^Δex3^ HCC, we performed CRISPR/Cas9 strategy *in vivo* to target the liver, using AAV8 hepatotropic vectors able to edit genes only in hepatocytes thanks to the hepatospecific TBG promoter driving the Cas9 expression (**Figure 2A**)^14^. As a proof of concept that such an approach can be used to induce long term carcinogenesis, we first generated the *Apc*^fs-ex15^ model. We selected specific guide RNA (sgRNA or sg) to target the beginning of the exon 15 of *Apc* (sg*Apc*) or the *Rosa26* locus (sg*Rosa*) as a control (**Figure 2B and figure S2A**). After production of rAAV8 viral particles, C57/Bl6 mice were injected with 2.10^11^ viral genome (vg) of AAV-CRISPR-sg*Apc*/*Rosa*. Two months after injection, editing efficiency on total liver was first quantified using TIDE algorithm (**Figure 2C-E-F**). Mean efficiency was 14.5±3.3% for sg*Apc* and 35.1±3.9% for sg*Rosa* showing a guide-dependent variability (**Figure 2C**). Despite this variability, the mutational profile induced by each guide as well as the total editing was highly reproducible (**Figure 2E-F**). We performed liver biopsies two weeks after injection of AAV-CRISPR-sg*Apc*. Single hepatocytes expressing GS were found randomly distributed in hepatic lobules, indicating focal activation of the β-catenin pathway (**Figure S3a,b**). At one month, these livers revealed a slightly higher editing efficiency (**Figure S3c,d**). As expected, control AAV-CRISPR-sg*Rosa*-injected mice had livers undistinguishable from wild type mice, as GS immunolocalization was restricted to the hepatocytes surrounding the central vein, which corresponds to the physiologic area with Wnt/βcatenin activation^11^ (Figure S4B).

**Figure 2:**
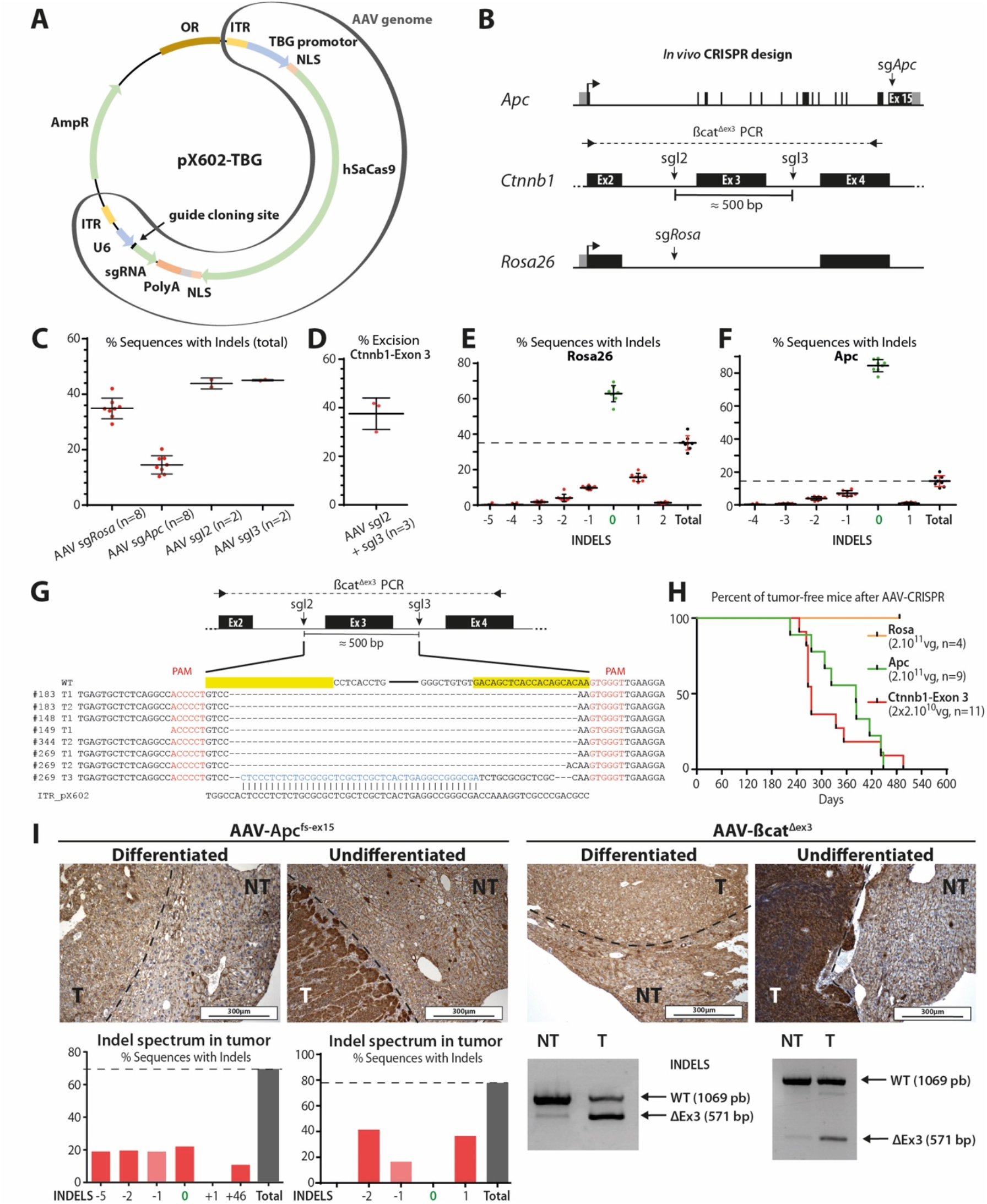
Tumor formation after AAV-CRISPR/Cas9 mediated loss-of-function of Apc or gain-of-function of β-catenin. **A:** pX602-TBG plasmid for the production of hepatotropic AAV8 vectors encoding the SaCas9 driven by a hepatospecific TBG promoter and a guide RNA. Small guides targeting *Apc*, *Ctnnb1*-Intron 2 (sgI2), *Ctnnb1*-Intron 3 (sgI3) or *Rosa26* locus (sgRosa) were each cloned into the plasmid. **B:** *In vivo* gene editing design. **C,D:** Total liver gene editing efficiency after AAV injection (2.10^11^vg total), assessed: (C) after 2 months via Tracking of indels by decomposition (TIDE); (D) after 1 month, via excision of *Ctnnb1*-Exon 3 (sgI2 + sgI3). **E-F:** Indel spectrum frequency on total liver 2 months post targeting of *Rosa26* or *Apc*. 0 indel corresponds to the WT sequence; −1 to −5 corresponds to a deletion of 1 to 5 nucleotides; +1 to +46 to an insertion of 1 to 46 nucleotides. **G:** *Ctnnb1*-Exon 3 editing in 8 tumors. **H:** kinetics of tumor detection by ultrasound. **I:** β-catenin staining of tumors induced via CRISPR-Apc^fs-ex15^ or CRISPR-βcat^Δex3^ (up) along with tumor editing (down) assessed via TIDE for Apc^fs-ex15^ and PCR for βcat^Δex3^.

Next, guides targeting intron 2 (sgI2) and intron 3 (sgI3) of *Ctnnb1* gene were selected to engineer a βcat^Δex3^ model (**Figure 2B, S2B-D**). Two months after injection of 2.10^11^ viral particles (vg) of single AAV-CRISPR-sgI2 or AAV-CRISPR-sgI3 in C57/Bl6 mice, editing efficiency was higher than 40% indels (**Figure 2C**). We then injected simultaneously 10^11^ vg of each AAV-CRISPR-sgI2 and AAV-CRISPR-sgI3. Such a dose induced β-catenin activation in a small subset of hepatocytes 2 weeks post injection to up to 40% of *Ctnnb1*-exon3 excision on total liver cells after 1 month (**Figure 2D, S4A)**. This led to a massive activation of the β-catenin pathway in more than 90% of hepatocytes and to the death of the injected animals. Lowering the dose to 2.10^10^ vg per AAV circumvented mortality. Resulting CRISPR-βcat^Δex3^ livers showed a significant number of single hepatocytes activated for β-catenin signaling, suitable for long term tumorigenesis studies (**Figure S4B**).

We performed a follow-up of CRISPR-*Apc*^fs-ex15^, CRISPR-βcat^Δex3^ and control CRISPR-*Rosa26* mice using ultrasound until tumor development (**Figure 2H**). As expected, all CRISPR-*Rosa26* mice remained healthy until sacrifice 16 months after injection (n=4) showing no deleterious effect of the CRISPR/Cas AAV-mediated *in vivo* strategy. On the contrary, all CRISPR-*Apc*^fs-ex15^ (n=9) and CRISPR-βcat^Δex3^ (n=11) mice developed tumors on average 355±73 days and 317±79 days after injection respectively. Editing analysis on 13 CRISPR-*Apc*^fs-ex15^ tumors using Tide revealed 1 to 4 indels per tumor, inducing frameshifts, with 75% of edited DNA in average, showing that tumors emerge from edited hepatocytes (**Table S3 and Figure 2I**). Analysis of exon 3 deletion in CRISPR-βcat^Δex3^ tumors and NT adjacent tissue showed a greater PCR amplification in tumors of the ~571 bp Δexon3 fragment expected (n=14) (**Figure 2I**). This was confirmed by Sanger sequencing of DNA in 8 tumors, which also revealed the insertion of a DNA sequence similar to the AAV-ITR in one tumor (**#269 T3 - Figure 2G**). Noteworthy, we targeted independently intron 2 or 3 to test the efficiency of the guides *in vivo* and no β-catenin activation was seen 2 months after injection (**Figure 2C, Figure S5A**). In this setting, one mouse that received a single AAV targeting intron 2 developed a β-catenin-activated tumor 8 months after injection (**Figure S5B**) and we found a deletion of more than 100bp at the beginning of exon 3. This oncogenic deletion is reminiscent of the heterozygous 45bp deletion that we identified in the murine Hepa 1-6 cell line (**Figure S5C**).

As described in CreLox-generated *Apc*^fs-ex15^ and βcat^Δex3^ mouse models, differentiated and undifferentiated tumors are observed when using CRISPR, showing an intense immunostaining for β-Catenin in UNDIFF tumors, and a slightly reinforced one in DIFF tumors (**Figure 2I**), indicating activation of the β-Catenin pathway. Our findings demonstrate that expressing a dominant stable mutant of β-Catenin is sufficient to induce tumorigenesis with a complete penetrance, and that CRISPR technology can easily and reproducibly be used to induce and maintain long term gene editing and carcinogenesis in the liver.

### βcat^Δex3^ mice develop tumors similar to *Apc*^fs-ex15^ mice

We analyzed mice with β-catenin-activated hepatic tumors, after the expression of a stable β-Catenin (βcat^Δex3^) or a loss-of-function of *Apc* (*Apc*^fs-ex15^), using either CreLox or CRISPR strategy. We could not detect lung metastases in either model, either macroscopically or by the analysis of hemalun-eosine stained FFPE-sections. We similarly found that *Apc*^fs-ex15^ and βcat^Δex3^ mice each gave rise to either DIFF or UNDIFF liver tumors (**Figure 3A, Figure S6**). Tumors were of similar size and were hyperproliferative (**Figure 3B,C**). Some tumor cells in UNDIFF tumors showed a positive immunostaining for the cleaved activated form of Caspase 3, revealing a mild apoptosis rate in these tumors (**Figure 3C**).

**Figure 3:**
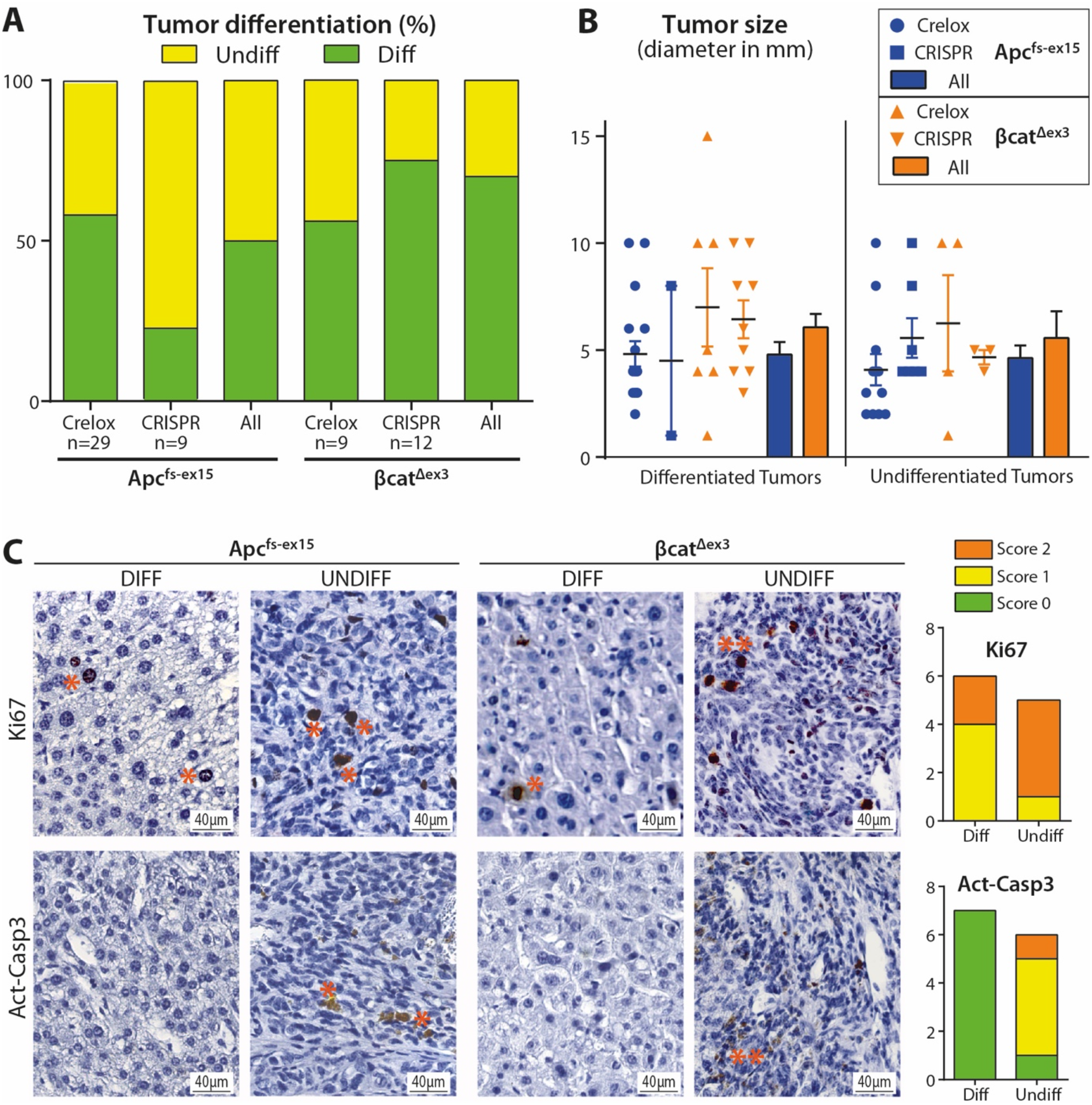
*Apc*^fs-ex15^ and βcat^Δex3^ tumors are phenotypically undistinguishable. **A:** Differentiation status in tumors from Crelox- and AAV-CRISPR-engineered *Apc*^fs-ex15^ and βcat^Δex3^ models. A chi-square test of independence shows no significant association between the genotype and differentiation status, **χ**^2^ (1, N = 60) = 3.34, p > 0,05. **B:** Tumor size is not statistically different between the models (One-way ANOVA). **C**: Proliferation (Ki67) and Apoptosis (Activated-Caspase 3) in tumors. Ki67 immunostaining reveals no statistically significant changes between differentiated and undifferentiated tumors, whereas activated Caspase 3 staining shows that a slight apoptosis can be detected only in undifferentiated tumors. Ki67 and Casp3 IHC were done on 7 diff. tumors from 5 *Apc*^fs-ex15^ (4 Crelox and 1 CRISPR) and 2 βcat^Δex3^ (Crelox) mice, and on 8 undiff. ones from 4 *Apc*^fs-ex15^ (2 Crelox and 2 CRISPR) and 4 βcat^Δex3^ (2 Crelox and 2 CRISPR) mice. No difference of staining was seen between tumors of *Apc*^fs-ex15^ versus βcat^Δex3^ genotypes. Scores for IHC were established as follows: Score 0 = no staining, score 1 = single tumor cells immunostained (*), score 2 = clusters of immunostained tumor cells (**).

We sought to examine the differences between *Apc*^fs-ex15^ and βcat^Δex3^ tumors from a transcriptomic point of view by RNA-sequencing of DIFF and UNDIFF tumors from both Crelox models. Strikingly, unbiased principal component analysis, hierarchical and consensus clusterings clearly separated into three distinct groups: NT tissue, DIFF tumors and UNDIFF tumors, independently on the model (**Figure 4**). Accordingly, almost no genes were found differentially expressed by comparing βcat^Δex3^ and *Apc*^fs-ex15^ tumors with similar phenotype (2 genes over-expressed in *Apc^f^*^s-ex15^ DIFF tumors compared to βcat^Δex3^ tumors and 0 genes between βcat^Δex3^ and *Apc*^fs-ex15^ UNDIFF tumors, q-value<0.05, −1>Log2(FC)>1) (**Figure S7**). Our data showed that tumors induced in Apc^fs-ex15^ model are similar to tumors induced in βcat^Δex3^ model.

**Figure 4:**
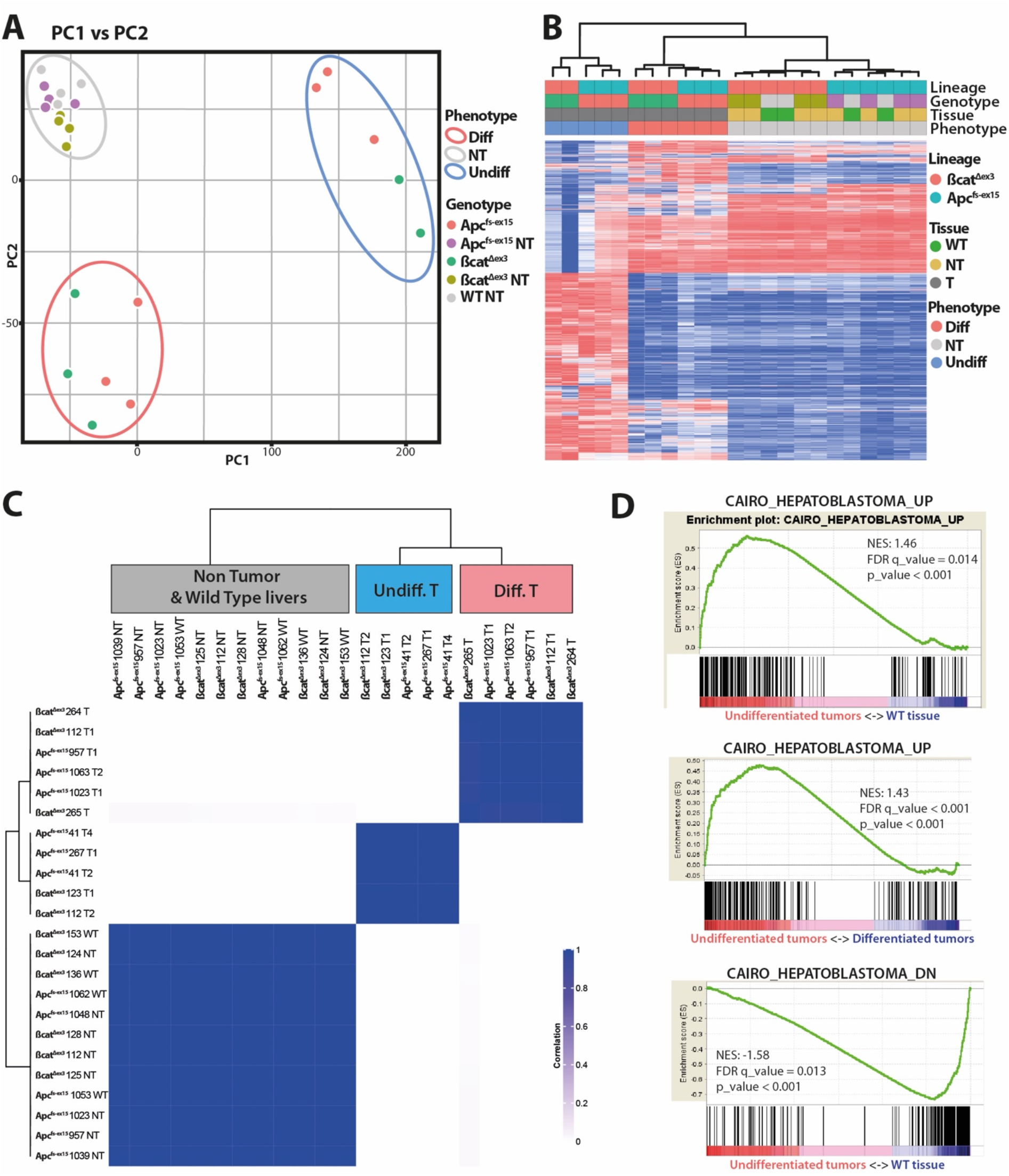
*Apc*^fs-ex15^ and βcat^Δex3^ tumors clusterize together, depending on their differentiation status. RNAseq data from non-tumoral (NT) wild type (WT) and tumoral tissues from Apc^fs-ex15^ and βcat^Δex3^ models using: **A:** Principal component analysis; **B:** unsupervised hierarchical clustering and **C:** consensus clustering (2000 genes). **D:** GSEA analysis showing an enrichment of a hepatoblastoma signature in mouse undifferentiated tumors compared to WT tissue or differentiated tumors.

### *Ctnnb1^ΔEx3^* and *Apc*^fs-ex15^ mouse models induce tumors transcriptionally close to human liver primary tumors

We previously described that DIFF mouse tumors looked like β-catenin-mutated human HCCs, described as well-differentiated tumors7. We searched to which type of liver primary tumors, UNDIFF βcatenin-activated mouse tumors could be related. A Geneset Enrichment Analysis (GSEA) found a human hepatoblastoma signature in these UNDIFF mouse liver tumors (**Figure 4D**).

We analyzed more deeply how murine hepatic tumors could be associated to subtypes of human liver primary cancers by doing an integrative analysis (**Figure S8**). We combined our recently characterized human hepatoblastoma (HB) cohort^16^ and HCC datasets from our previously described molecular classification G1 to G6^17, 18^, with RNAseq data of mouse βcat-HCCs. Using hierarchical clustering, we clearly showed that: (1) DIFF mouse tumors are close to human well-differentiated β-Catenin-activated G5-G6 HCCs (βcat-hHCC) constituting the cluster 4; (2) UNDIFF mouse tumors are close to human hepatoblastomas, particularly to mesenchymal hepatoblastomas (M-hHB), constituting the cluster 1 (**Figure 5**). A similar clustering was obtained when generating a correlation matrix between human and mouse samples (**Figure S9A**). As expected from their β-catenin-activated status, both UNDIFF and DIFF mouse HCCs had a β-catenin transcriptional signature, but this signature was qualitatively different. The DIFF tumors were enriched in β-catenin signaling targets found in the adult liver after Apc loss, expressing pericentral markers such as GS^3^ (**Figure S9B**). This pericentral β-catenin signature is also found in human βcat-hHCCs^3^. Conversely, UNDIFF tumors were enriched in the expression of β-catenin targets found in the embryonic liver after Apc loss^19^ (**Figure S9B**): some of which are found common in M-hHB, such as NKD1, TNFRSF19, APCDD1 (**Figure 5**). Besides this common β-catenin progenitor signature, we found a common Epithelial-Mesenchymal Transition (EMT) signature in UNDIFF tumors and M-hHBs (**Figure 6B**). This included an increase in mesenchymal markers such as MMP2 and Vimentin (VIM), an increase in transcription factors involved in EMT, such as SNAIL and SLUG (SNAI2), ZEB1 and ZEB2, together with a decrease in epithelial markers such as Occludin (OCLN) and E-cadherin (CDH1) (**Figure 6B**). We confirmed this pattern, as we found a strong Vimentin immunostaining only in UNDIFF tumors (**Figure 6C**). This underlines the mesenchymal status of murine UNDIFF tumors.

**Figure 5:**
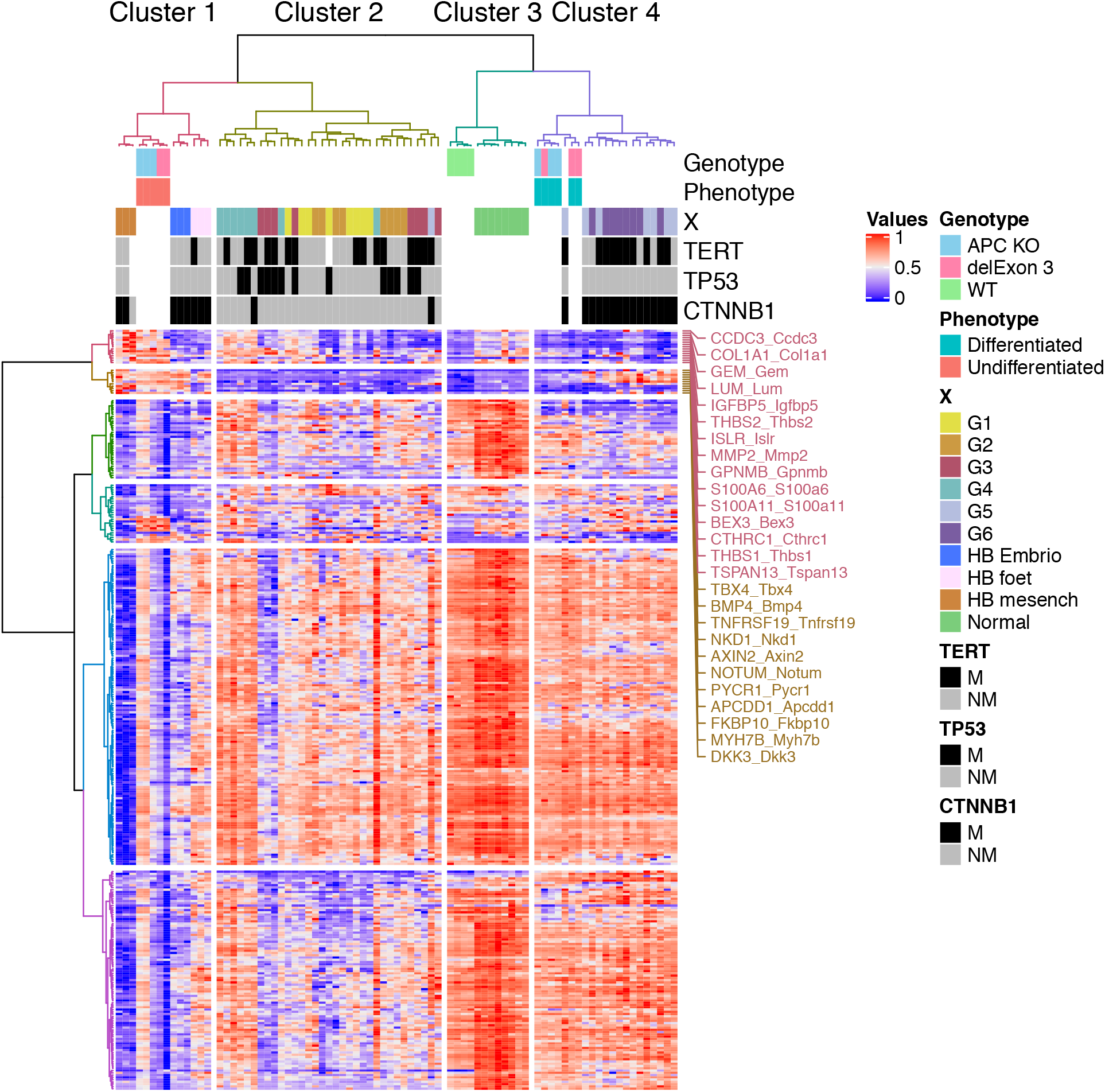
Hierarchical clustering of the integrated human/mouse analysis. The 60 samples are separated into 4 clusters. Cluster 1: Human hepatoblastoma (HB) samples and undifferentiated mouse tumors. Clusters 2: G1 to G4 HCCs. Cluster 3: Human and mouse non-tumoral tissues. Cluster 4: G5 and G6 HCC and differentiated mouse tumors. M, Mutated; NM; non-mutated.

**Figure 6:**
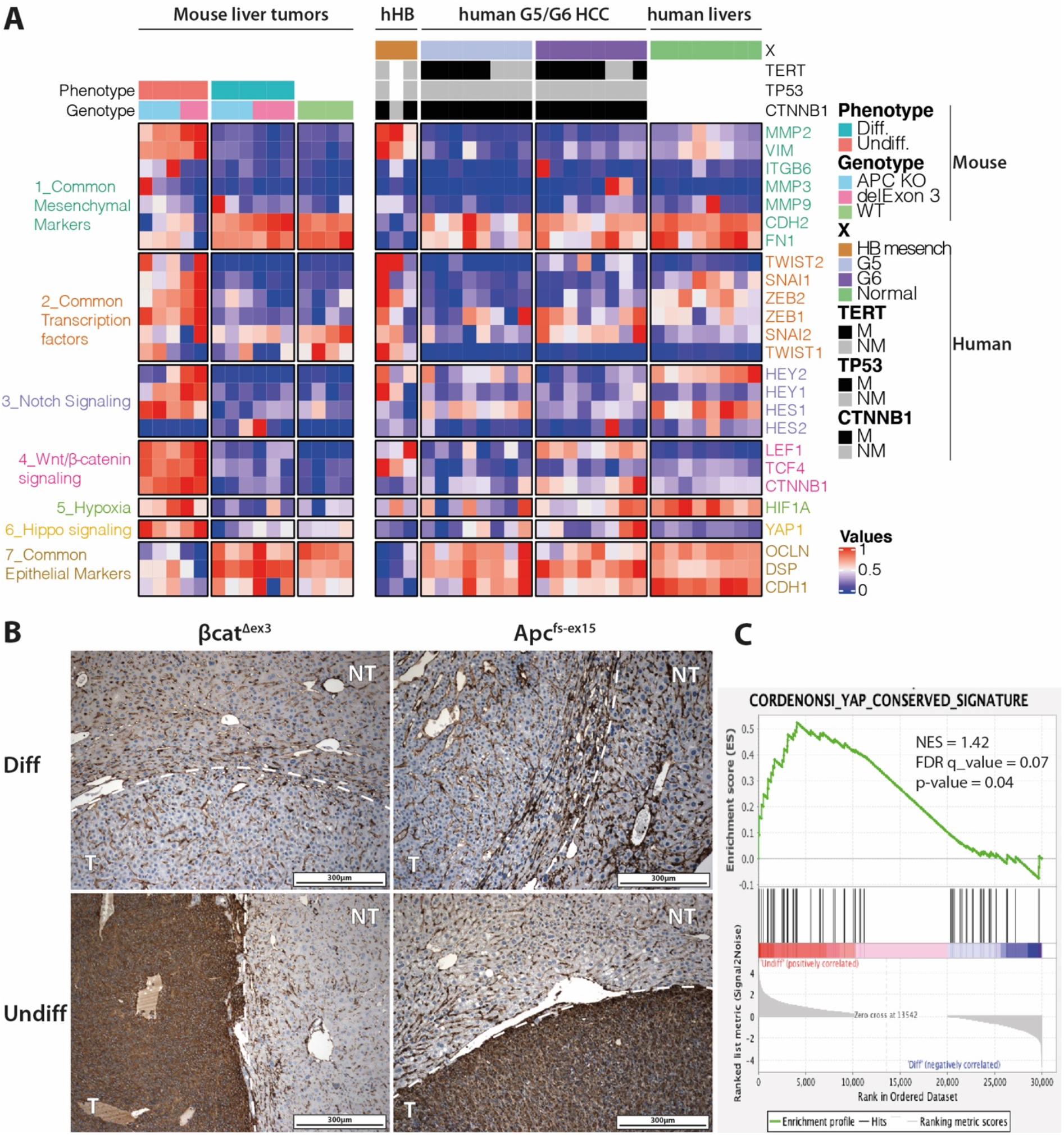
Mesenchymal and YAP signature in undifferentiated β-catenin-activated mouse HCCs. **A.** Expression profile of common EMT related markers in mouse and human samples (Mesenchymal HB, G5-G6 HCCs). **B.** Vimentin IHC performed on DIFF and UNDIFF mouse tumors from *Apc*^fs-ex15^ and *βcat*^Δex3^ models. **C.** GSEA showing an enrichment in YAP conserved signature in UNDIFF compared to DIFF mouse tumors.

To summarize, *Apc*^fs-ex15^ and βcat^Δex3^ mouse tumors similarly generate two distinct types of tumors transcriptionally close to human β-catenin-activated tumors, either well-differentiated G5-G6 HCCs, or mesenchymal HBs.

### YAP/TAZ signaling is activated in undifferentiated β-catenin-activated mouse liver tumors

We finally looked for a pathway directing the phenotypic features of UNDIFF mouse tumors, and connected to β-catenin pathway in Hepatoblastoma. The YAP/TAZ pathway was a tempting hypothesis, as it is activated in hepatoblastomas and linked to β-catenin^20^, ^21^. Accordingly, a GSEA showed an enrichment in the Cordenonsi YAP Conserved Signature consisting in target genes of YAP/TAZ signaling^22^ (**Figure 6D, Figure S10A**). The partners of YAP/TAZ signaling Yap1, Taz (Wwtr1) and the transcription factors involved in Yap/Taz-dependent gene expression: Tead1,2,3,4, were also overexpressed in UNDIFF tumors, **(Figure S10B)**. We thus characterized this pathway *in situ* in control livers (**Figure 7, Figure S11**), in DIFF and UNDIFF tumors by IHC (**Figure 7**). We found a zonal expression of Taz protein in pericentral hepatocytes in control livers. Taz was abundant in DIFF tumors but absent from UNDIFF ones. Tead1 protein was expressed in bile ducts of control livers and at the periphery of DIFF tumors, but was absent from DIFF tumor cells. Conversely, UNDIFF tumors strongly expressed Yap1 and TEAD1, with no expression of Taz. By combining the data showing the overexpression of Yap target genes in UNDIFF tumors with those showing the colocalization of Yap1 and Tead1 in the nucleus of UNDIFF tumor cells, we concluded that a Yap/Tead1 signaling cascade promotes the expression of oncogenic Yap targets in UNDIFF mouse tumors.

**Figure 7:**
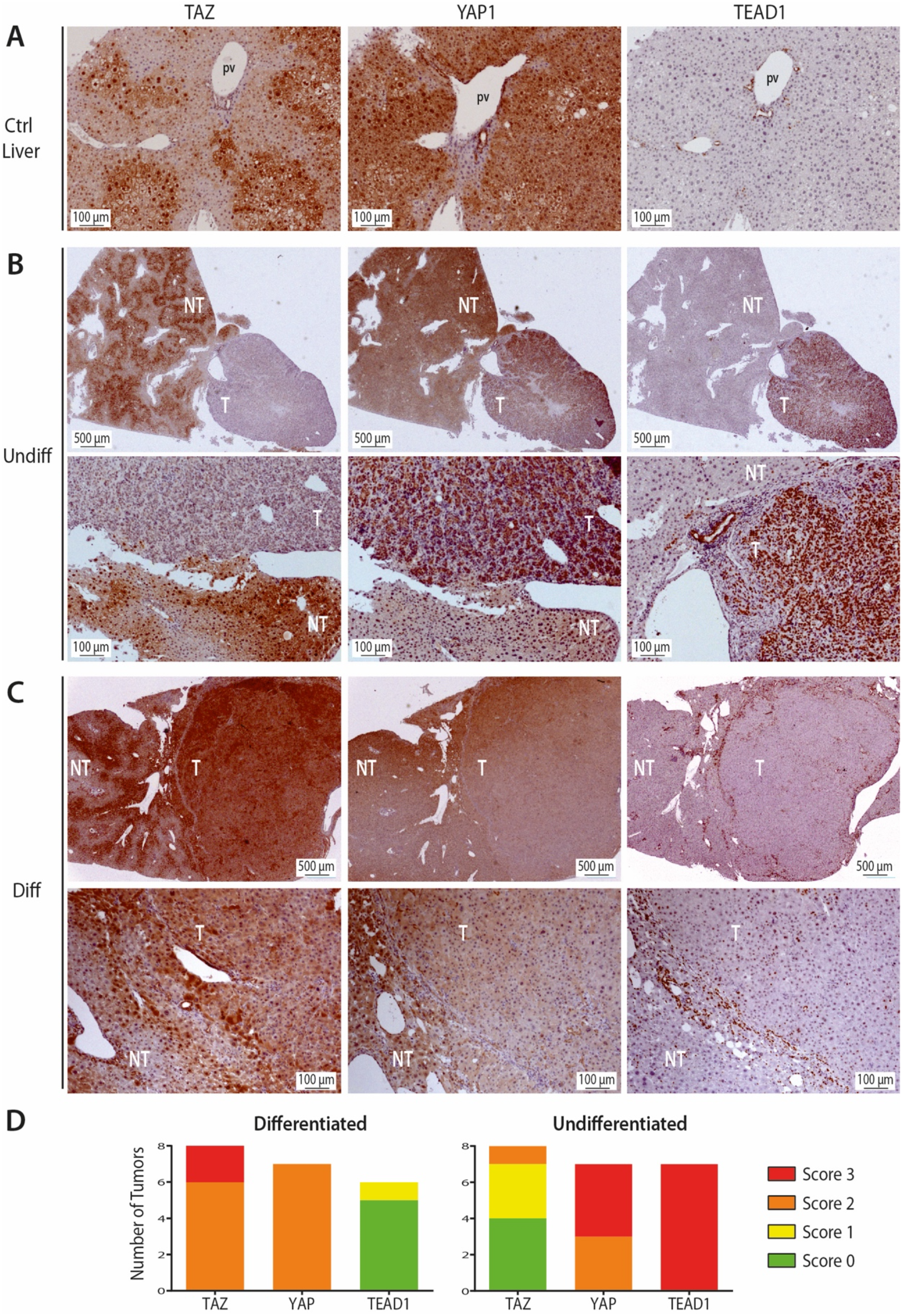
YAP/TAZ pathway is distinctly activated in undifferentiated versus differentiated mouse HCCs. Immunostainings for Taz, Yap1 and Tead1 in representative sections of : **A.** control liver, **B.** UNDIFF mouse tumor, **C.** DIFF mouse tumor. Pv= portal vein; T=Tumor; NT= Non Tumoral adjacent liver. **D.** Immunoscorings. Score 0 = no staining in T, score 1 = staining in T < NT, score 2 = staining in T = NT, score 3 = nuclear staining in T > NT. DIFF tumors were from 5 *Apc*^fs-ex15^ (3 Crelox and 2 CRISPR) and 2 βcat^Δex3^ (1 CRISPR and 2 Crelox) mice. UNDIFF ones came from 5 *Apc*^fs-ex15^ (2 Crelox and 3 CRISPR) and 3 βcat^Δex3^ (2 Crelox and 1 CRISPR) mice.

## Discussion

Mouse models of β-catenin dependent tumorigenesis have been widely used. However, while HCC development can be induced by bi-allelic loss-of-function of *Apc* in mice, it was believed since 2002 that the expression of a dominant stable mutant of β-catenin is not sufficient to induce tumorigenesis^10^. It was supposed that liver carcinogenesis requires additional oncogenic events to emerge, justifying the development of new models of β-catenin-activated oncogenesis, in which the gain-of-function of β-catenin was combined to that of H-Ras^23^, c-Met^24^, Akt^25^ or YAP^21^. These models brought significant information concerning the oncogenic cooperations involved with β-catenin signaling, which speed up tumor emergence, and emphasized the fact that β-catenin activation is not oncogenic *per se* in the liver. Moreover, this observation raised interrogations regarding the relevance of mouse models using *Apc* loss-of-function.

Here we first showed that β*cat*^Δex3^ mouse model can actually induce tumorigenesis without engineering additional genetic alterations in mice, suggesting that β-catenin activation is an initiating event for liver carcinogenesis, even if a latency period of 6 to 9 months is observed before tumor detection. This latency period can be due to the fact that activating β-catenin signaling does not promote proliferation in single hepatocytes that remain quiescent, but changes their metabolic and epigenetic properties^13, 26, 27^. The oncogenicity of β-catenin signaling in the liver is thus a longstanding process, which remains to be elucidated^28^. The latency between β-catenin activation and tumorigenesis can by itself explain why a dominant stable β-catenin was previously described as not tumorigenic in the liver, due to a 6-month tumor follow-up^10^. On the other hand, as for the *Apc* model, the number of βcat^Δex3^ hepatocytes needs to reach a threshold above which tumor development can be observed during the animal lifetime, while being low enough not to impair liver functions with a subsequent lethality. Interestingly, no mortality, full penetrance and a high reproducibility was observed using CRISPR-mediated exon 3 deletion, showing that this AAV-CRISPR approach is the method of choice to generate β-catenin-activated liver tumors.

Two types of tumors, discriminated by their phenotype (differentiated or undifferentiated), were generated using *Apc^fs-ex15^* or βcat^Δex3^ models. Transcriptomic and phenotypic analysis clearly show that generated tumors are indistinguishable depending on the model used but are strongly clustering depending on the differentiation status. This first supports the hypothesis that *Apc* loss-of-function drives hepatocarcinogenesis through the activation of β-catenin signaling, meaning that the alternative pathways in which Apc is involved, such as chromosomal instability promoted by Apc loss in colorectal cancers, does not prevail in HCC initiation^8^. Second, our data show two distinct oncogenesis pathways driven by β-catenin signaling in the liver. Integrated analysis of murine samples with human HCC and hepatoblastomas indicates that mouse differentiated tumors are transcriptionally close to G5-G6 human HCCs which are well differentiated and mutated for *CTNNB1*. Murine undifferentiated tumors cluster with human hepatoblastomas mostly mutated in *CTNNB1*. Interestingly, the three mesenchymal human hepatoblastoma, two of which being *CTNNB1* mutated, are closer to murine undifferentiated tumors. The origin of hepatoblastoma-like tumors in our models is unknown. The development of HCC and hepatoblastoma following expression in fetal liver progenitor cells of a stable mutant of β-catenin has been already described in mice, raising the possibility that undifferentiated tumors arise from immature/progenitor cells in the liver^29^. This would suggest that in our study, liver progenitors could be targeted by the Cre-lox system or AAV vectors to deliver the CRISPR machinery. However, AAV8 vectors coupled with TBG promoter have been described highly specific in targeting hepatocytes without targeting any other hepatic compartment^25^. Moreover, the TTR-Cre^Tam^ mouse used for CreLox strategy was described to be active both in the fetal and the adult liver, but we could show β-catenin accumulation only in hepatocytes, after Tamoxifen injection to adult mice^13,14^. In agreement with this, previous studies support the fact that HCC originates from hepatocytes and not from progenitor/biliary compartment^30^. Lastly more and more studies suggest that hepatocytes and cholangiocytes are the main sources of hepatic progenitors in the liver^31–33^, and this suggests that both differentiated and undifferentiated tumor types arise from mature hepatocytes. We hypothesize that different additional oncogenic events elicited by β-catenin activation in hepatocytes, could dictate the path toward a more or less differentiated cell fate and lead to specific tumor phenotypes.

Our data also strengthen the hypothesis that distinct Yap/Taz signalings could be players of either differentiated HCCs or poorly differentiation liver tumors. Yap and Taz are not identical twins, as recently reviewed in other biological systems^34^. In the liver, one cascade could be dictated by Taz together with β-catenin: Taz expression is limited to β-catenin-activated pericentral hepatocytes in control livers, and it is found together with Yap1 in β-catenin-activated DIFF mouse tumors. This suggests that Taz and β-catenin could be interactive inducers of pericentral gene expression, both in pericentral hepatocytes, and in DIFF mouse tumors and human G5/G6 HCCs^3^. On the other hand, Yap1 has been described as overriding the pericentral zonation program and directing periportal hepatocytes to dedifferentiate into atypical ductal cells^35^. Accordingly, we found here that immunostainings for both Yap1 and its nuclear effector Tead1 are more intense in and around the bile ducts. They are co-expressed in β-catenin-activated UNDIFF tumors, which exhibit small and basophilic tumor cells with atypical ductal cells features. This strongly suggests a Yap1/Tead1 cascade in UNDIFF tumors, and potentially in Hepatoblastomas.

Lastly, we described the possibility to use CRISPR-mediated gene editing for studying long term tumorigenesis. Some concerns have been raised regarding the persistence of edited hepatocytes due to the immunogenic properties of the SaCas9. While a recent study describes the loss of edited hepatocytes with time in a model of SaCas9 pre-immunized mice, we did not observe such a decrease^36^. Another concern regarding *in vivo* editing using CRISPR is the specificity of the edition. We found an unwanted outcome, as the sequencing of one βcat^Δex3^ tumor revealed the insertion of 42bp similar to the AAV ITR. Insertion of AAV have been: (1) described as potential oncogenic events in human HCC^37, 38^; (2) described as common at CRISPR-induced Double-Stranded Breaks^39^. A retrospective study on the use of AAV vector in therapy (not combined with CRISPR) clearly described AAV vectors as safe in clinical settings^40^. Unwanted tumorigenesis due to AAV insertion in mouse experimental settings have not been reported yet. Altogether the easy setting, and time saving aspect of CRISPR mediated *in vivo* gene editing compared to the development of new Cre-lox models makes it very attractive to study liver carcinogenesis^41^.

## Supporting information

Supplemental_Data_Loesch

## Acknowledgements

We thank Pr Feng Zhang for the gift of AAV8-CRISPR-SaCas9 plasmids, and Dr Violaine Moreau for helpful discussions. We are grateful to the animal facilities of Cochin Institute and of the Centre de Recherche des Cordeliers for mice care, and to the Live Imaging Platform of Cochin Institute for ultrasound follow-up. We also thank Yoan Renoux-Martin for Yap, Taz and Tead1 immunostainings.

