## Supplemental_Data_Loesch for "Deleting *in vivo* β-catenin degradation domain in mouse hepatocytes drives hepatocellular carcinoma or hepatoblastoma-like tumors"

**Sequence guides used for targeting specific loci**

| Target | sgRNA / PAM used in vitro | Strand | Length |
|---|---|---|---|
| *Ctnnb1-I2-1* | 5'-AAATAATCAGCAAGCCACCGATGGGAT-3' | + | 21 |
| *Ctnnb1-I2-2* | 5'-GCAAGCCACCGATGGGATCTAATGAGT-3' | + | 21 |
| ***Ctnnb1-I2-3*** | **5'-TCAGTATGAGCTCCATGGGACAGGGGT-3'** | **-** | **21** |
| ***Ctnnb1-I3-1*** | **5'-GACAGCTCAGCCACAGCACAAGTGGGT-3'** | **+** | **21** |
| *Ctnnb1-I3-2* | 5'-AAGTGGGTTGAAGGAAGGGCGGAGGGT-3' | + | 21 |
| *Ctnnb1-I3-3* | 5'-TGAAGGAAGGGCGGAGGGTAGCGGAGT-3' | + | 21 |
| *Apc-1* | 5'-AATAAGGCTGACATCTGTGCTGTGGATG-3' | + | 21 |
| *Apc-2* | 5'-AGCCATTATTGAAAGTGGAGGTGGGAT-3' | + | 21 |
| ***Apc-3*** | **5'-TCTTCAGAATAGGATTCAACTGAGGGT-3'** | **-** | **21** |
| ***Rosa-1*** | **5'-CTCGATGGAAAATACTCCGAGGCGGAT-3'** | **-** | **21** |
| *Rosa-2* | 5'-GTAGATTAAAGACATGCTCACCCGAGT-3' | + | 21 |
| *Rosa-3* | 5'-TCAAGCAGGAGAGTATAAAACTCGGGT-3' | - | 21 |
| **Target** | **sgRNA / PAM used in vivo** | **Strand** | **Length** |
| *Rosa26 (Rosa-1)* | **5'-CTCGATGGAAAATACTCCGAGGCGGAT-3'** | - | 21 |
| *Apc (Apc-3)* | **5'-TCTTCAGAATAGGATTCAACTGAGGGT-3'** | - | 21 |
| *Ctnnb1-I2 (I2-3)* | **5'-TCAGTATGAGCTCCATGGGACAGGGGT-3'** | - | 21 |
| *Ctnnb1-I3 (I3-1)* | **5'-GACAGCTCAGCCACAGCACAAGTGGGT-3'** | + | 21 |

**Primers for sequencing, for TIDE and excision analyses**

| Target | PCR primers used in vitro | | | Sequencing |
|---|---|---|---|---|
| **PCR** | **Forward** | **Reverse** | **Size** | **for TIDE** |
| *Rosa26* | 5'-CTTGCTCTCCCAAAGTCGCT-3' | 5'-CCAATGCTCTGTCTAGGGGT-3' | 709 | Reverse |
| *Ctnnb1-I2* | 5'-TTTTGGTGTCGGGGCACATA-3' | 5'-TTGCTCTTGCGTGAAGGACT-3' | 669 | Forward |
| *Ctnnb1-I3* | 5'-CTGGCAGCAGCAGTCTTACT-3' | 5'-CATGGTGCGTACAATGGCAG-3' | 798 | Forward |
| *Ctnnb1-Ex3* | 5'-TTTTGGTGTCGGGGCACATA-3' | 5'-CATGGTGCGTACAATGGCAG-3' | 1069 | |
| Apc-1 & Apc-2 | 5′-CCTGTGTTGACTCATAGAAACAGC-3′ | 5′-GCATGGCTGGATTTCTCAACTAC-3′ | 616 | Forward |
| Apc-3 | 5'-AGTTCTGCTTCTACCACCGAG-3' | 5'-TCCTGACACAGAGACTGGTTTAC-3' | 683 | Forward |
| **Target** | **PCR primers used in vivo** | | | **Sequencing** |
| **PCR** | **Forward** | **Reverse** | **Size** | **for TIDE** |
| *Rosa26* | 5'-CTTGCTCTCCCAAAGTCGCT-3' | 5'-CCAATGCTCTGTCTAGGGGT-3' | 709 | Reverse |
| *Apc (Apc-3)* | 5'-AGTTCTGCTTCTACCACCGAG-3' | 5'-TCCTGACACAGAGACTGGTTTAC-3' | 683 | Forward |
| *Ctnnb1-I2* | 5'-TTTTGGTGTCGGGGCACATA-3' | 5'-TTGCTCTTGCGTGAAGGACT-3' | 669 | Forward |
| *Ctnnb1-I3* | 5'-CTGGCAGCAGCAGTCTTACT-3' | 5'-CATGGTGCGTACAATGGCAG-3' | 798 | Forward |
| *Ctnnb1-Ex3* | 5'-TTTTGGTGTCGGGGCACATA-3' | 5'-CATGGTGCGTACAATGGCAG-3' | 1069 | |

**Table S1:** Oligonucleotides used as sequence-guide primers, and for PCR amplification followed by Sanger sequencing and TIDE analysis or excision assessment (βcat-exon 3).

| Antibodies against | Description | | Supplier | Reference | Dilution |
|---|---|---|---|---|---|
| GS | Mouse, monoclonal | Primary | BD Biosciences | 610518 | 1/400 |
| β-catenin | Mouse, monoclonal | Primary | BD Biosciences | 610153 | 1/50 |
| Yap1 | Rabbit, monoclonal | Primary | Cell Signaling | D8H1X | 1/500 |
| Taz (Wwtr1) | Rabbit, monoclonal | Primary | Cell Signaling | E9J5A | 1/200 |
| Tead1 | Rabbit, monoclonal | Primary | Cell Signaling | D9X2L | 1/50 |
| Ki67 | Rabbit, monoclonal | Primary | Abcam | Ab16667 | 1/150 |
| Cleaved-Casp3 | Rabbit, polyclonal | Primary | Cell Signaling | 9661 | 1/600 |
| Vimentin | Rabbit, polyclonal | Primary | Abcam | Ab137321 | 1/100 |
| **Secondary antibodies** | | | | | |
| M.O.M., mouse on mouse | Anti-mouse, biotin | Secondary + background elimination | Vector Lab | BMK2202 | Kit |
| Histofine | Anti-rabbit, HRP | Secondary | Microm | F/414341F | Kit |

**Table S2**: Antibodies used for Immunohistochemistry.

| | % Indels | | | | | | | | | | | | | | |
|---|---|---|---|---|---|---|---|---|---|---|---|---|---|---|---|
| Mouse | **Total** | -40 | -38 | -29 | -8 | -5 | -4 | -2 | -1 | 0 | +1 | +2 | +8 | +10 | +46 |
| 101 T1 | **70.7** | | | | | | | 70.5 | | 27.4 | | | | | |
| 101 T2 | **81.9** | | | | | | | | 76.1 | 12.3 | | | | | |
| 102 T1 | **69.8** | | | | | | 16.7 | | 51.8 | 26.8 | | | | | |
| 103 T1 | **30.7** | | 1.2 | 5.8 | 13.8 | | | | 6.4 | 66.5 | | | | | |
| 138 T1 | **84.7** | | | | | | | | 64.7 | 13.73 | 19.5 | | | | |
| 139 T1 | **61.5** | | | | | | | 21.2 | 22.1 | 34.1 | 18.2 | | | | |
| 139 T2 | **69.1** | | | | | 19 | | 20 | 19.2 | 22.3 | | | | | 10.9 |
| 139 T3 | **79.1** | | | | | | | | 42 | 17.3 | | 37.1 | | | |
| 141 T1 | **61.7** | | | | | | | 16.2 | 35 | 30.6 | | | 10.5 | | |
| 141 T2 | **82.5** | | | | | | | | 80.9 | 16.2 | | | | | |
| 262 T1 | **59.2** | 14.6 | | | | | | | 20.2 | 34.7 | 9.2 | | | 15.1 | |
| 264 T1 | **81.9** | | | | | | | 41.6 | 40.3 | 14.4 | | | | | |
| 342 T1 | **72.4** | | | | | | | | 70.7 | 25.1 | | 1.1 | | | |

**Table S3: Indel spectrum in each tumor developed after loss of function of *Apc* using CRISPR *in vivo*.** *P*-value < 0.001 for each estimated indel abundance. Green : Estimated percentage of WT sequence. Gray : Estimated percentage of indel type. 0 indel corresponds to the WT sequence ; -1 to -40 corresponds to a deletion of 1 to 40 nucleotides ; +1 to +46 to an insertion of 1 to 46 nucleotides.

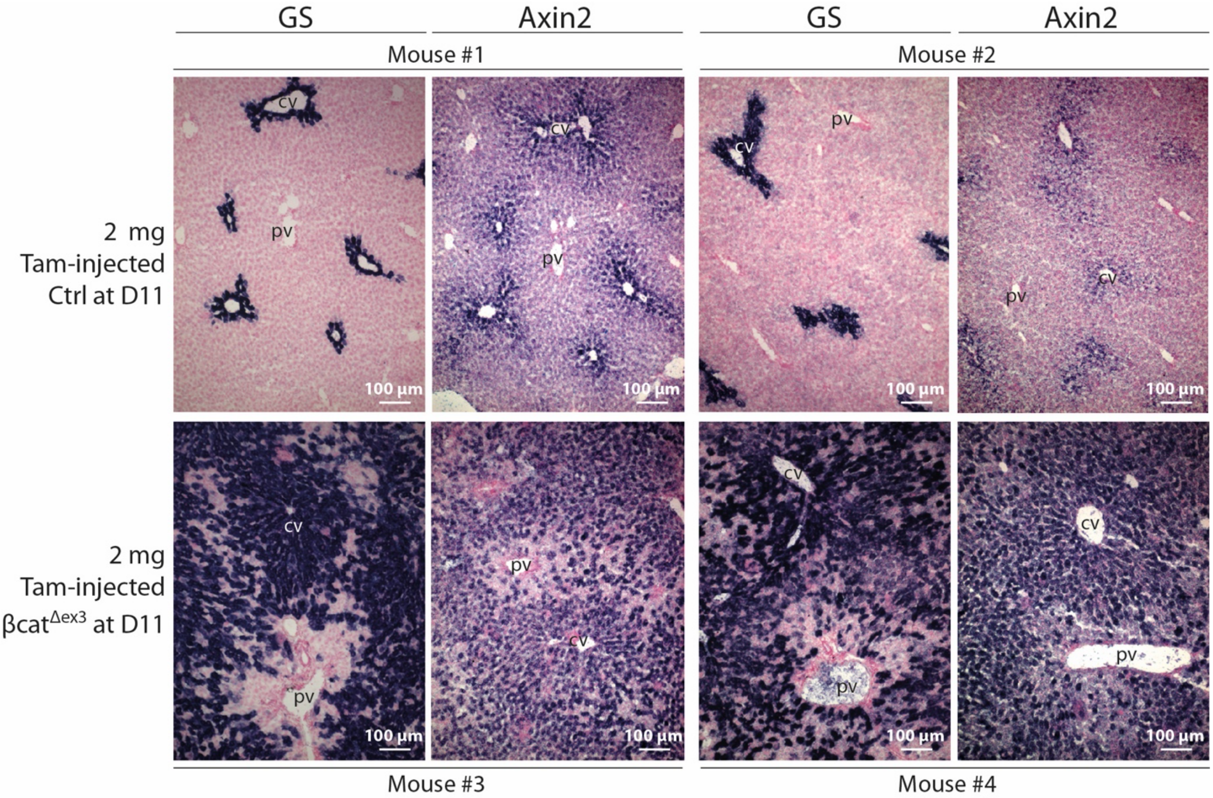


**Figure S1: *In situ* hybridization of Axin2 and Glutamine Synthetase (GS) mRNAs, two β-catenin hepatospecific targets in Cre-loxP-engineered βcat$^{\Delta ex3}$ livers after high dose of Tamoxifen injection.** Tamoxifen (2 mg) was intraperitoneally injected to compound βcat$^{ex3\text{-}flox}$/TTR-Cre$^{Tam}$ mice (βcat$^{\Delta ex3}$) and to βcat$^{ex3\text{-}flox}$ mice as controls. Note the physiological staining for Axin2 and GS due to Wnt signaling in the pericentral (PC) hepatocytes surrounding the central vein (cv), and its absence in periportal (PP) hepatocytes surrounding the portal vein (pv). In βcat$^{\Delta ex3}$ livers, Axin2 and GS stain all the hepatocytes at the exception of the most PP ones. Two mice of each genotype were analyzed (Ctrl: #1 & #2, βcat$^{\Delta ex3}$: #3 & #4). *In situ* hybridizations were performed with the digoxygenin probes and according to the protocol described in (Benhamouche et al., Dev Cell 2006).

A

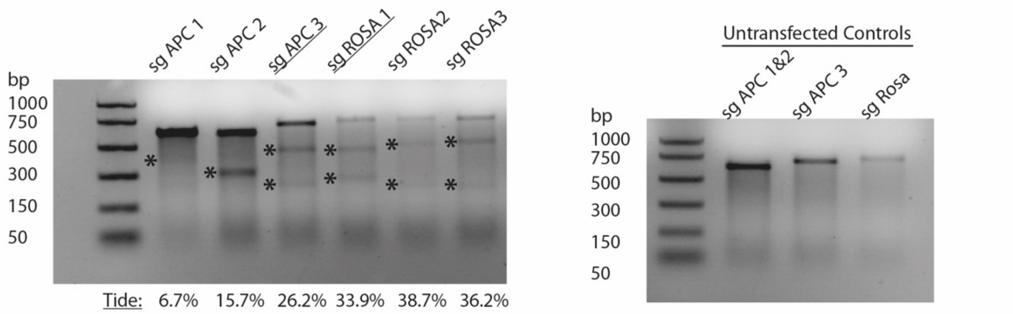


B

| Guide RNA for *Ctnnb1* | Promoter CMV - Indels in vitro | Promoter TBG - Indels in vitro |
|---|---|---|
| sgI2-1 : 5' AAATAATCAGCAAGCCACCGATGGGAT 3' | 14.1 % | 8.4 % |
| sgI2-2 : 5' GCAAGCCACCGATGGGATCTAATGAGT 3' | 9.3 % | 7.1 % |
| sgI2-3 : 5'TCAGTATGAGCTCCATGGGACAGGGGT 3' | 22.6 % | 11.2 % |
| sgI3-1 : 5' GACAGCTCAGCCACAGCACAAGTGGGT 3' | 18.3 % | 10.8 % |
| sgI3-2 : 5' AAGTGGGTTGAAGGAAGGGCGGAGGGT 3' | 3 % | 2.1 % |
| sgI3-3 : 5'TGAAGGAAGGGCGGAGGGTAGCGGAGT 3' | 10.7 % | 4.4 % |

C

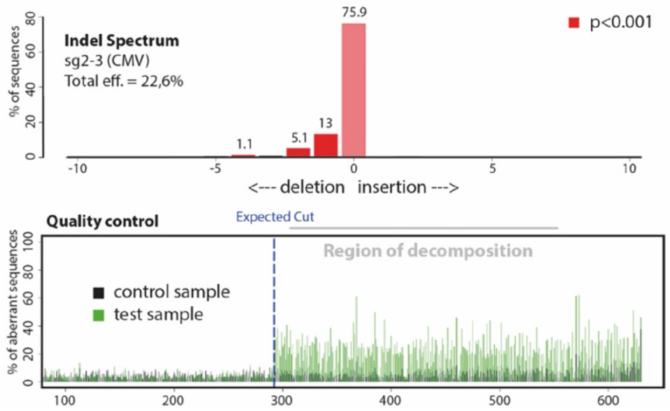


D

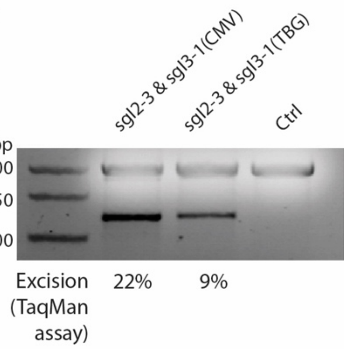


**Figure S2: Guide selection in vitro. A :** T7 nuclease assay and Tide results after pX601 (Promotor CMV) transfection in Hepa 1-6 mouse hepatoma cells to test the editing efficiency of 6 different guides targeting *Rosa26* or *Apc* (exon 14 for sgAPC 1 and 2 or exon 16 for sgAPC 3). The asterisks depict the the edited amplicon. sgAPC3 and sgROSA1 have been selected for injecting in vivo AAV-CRISPR-Cas9 particles. The controls correspond to a T7 nuclease assay performed on untransfected cells. Hepa1-6 cells were chosen because they correctly expressed wild-type Apc. As they have an activating truncation in Ctnnb1-exon3 (see Figure S5C) and are strongly β-catenin-activated, we did not use this cell line for testing the efficacy of Ctnnb1-exon3 deletion. **B :** Tide results after pX601 (promoter CMV) or pX602 (Promoter TBG) transfection in AML12 mouse hepatic cells to test the editing efficiency of 6 different guides targeting *Ctnnb1* intron 2 or 3. **C :** example of tide analysis for sg2-3 CMV. **D :** Deletion of exon 3 was then assessed on gel (down) after co-transfection of the guides selected (underlined in the Table S2B) in AML12 using pX601 or pX602 (Ctrl : untransfected cells).

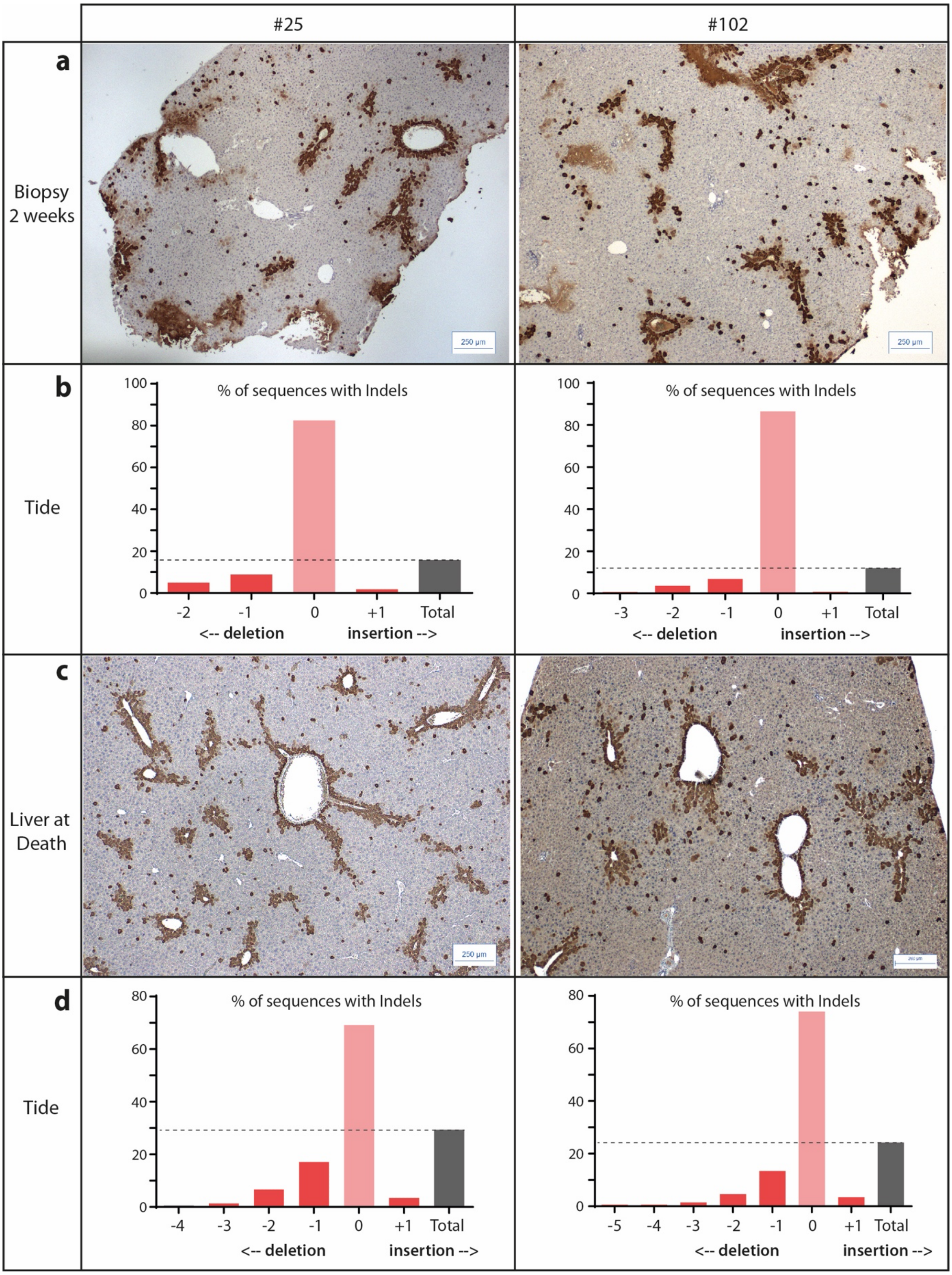


**Figure S3**: **Long-term stability of Apc$^{fs\text{-}ex15}$ editing in mouse livers. (a,c)** Glutamine synthetase immunostaining and **(b,d)** indel spectrum analysis using Tide. Two mice were analyzed post injection of AAV-CRISPR-Apc$^{fs\text{-}ex15}$, **(a,b)** 2 weeks after biopsy, or **(c,d)** at death (8 and 14 months for #25 and #102 respectively).

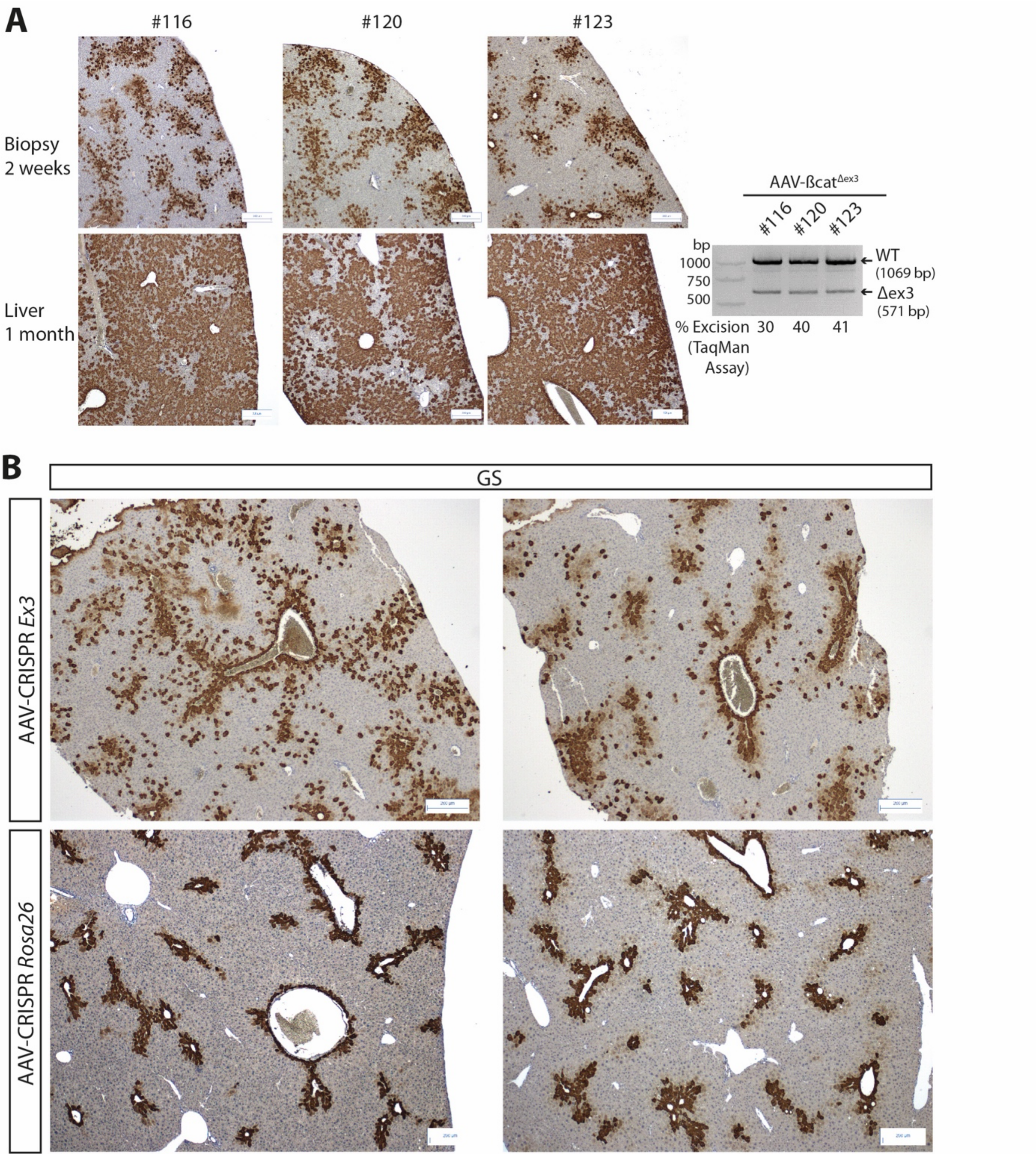


**Figure S4: AAV-CRISPR-engineered βcat$^{\Delta ex3}$ livers.** GS immunostaining (IHC) on liver: **A:** two weeks and one month after injection of $10^{11}$ vg of each AAV-CRISPR-sgI2 (sgI2) and AAV-CRISPR-sgI3 (sgI3) along with the corresponding PCR surrounding exon 3 (ex3-PCR); **B:** one month after the injection of $2.10^{10}$ vg of each AAV-CRISPR-sgI2 and AAV-CRISPR-sgI3. Control mice are those injected with $2.10^{11}$ vg of AAV-CRISPR-*Rosa26*.

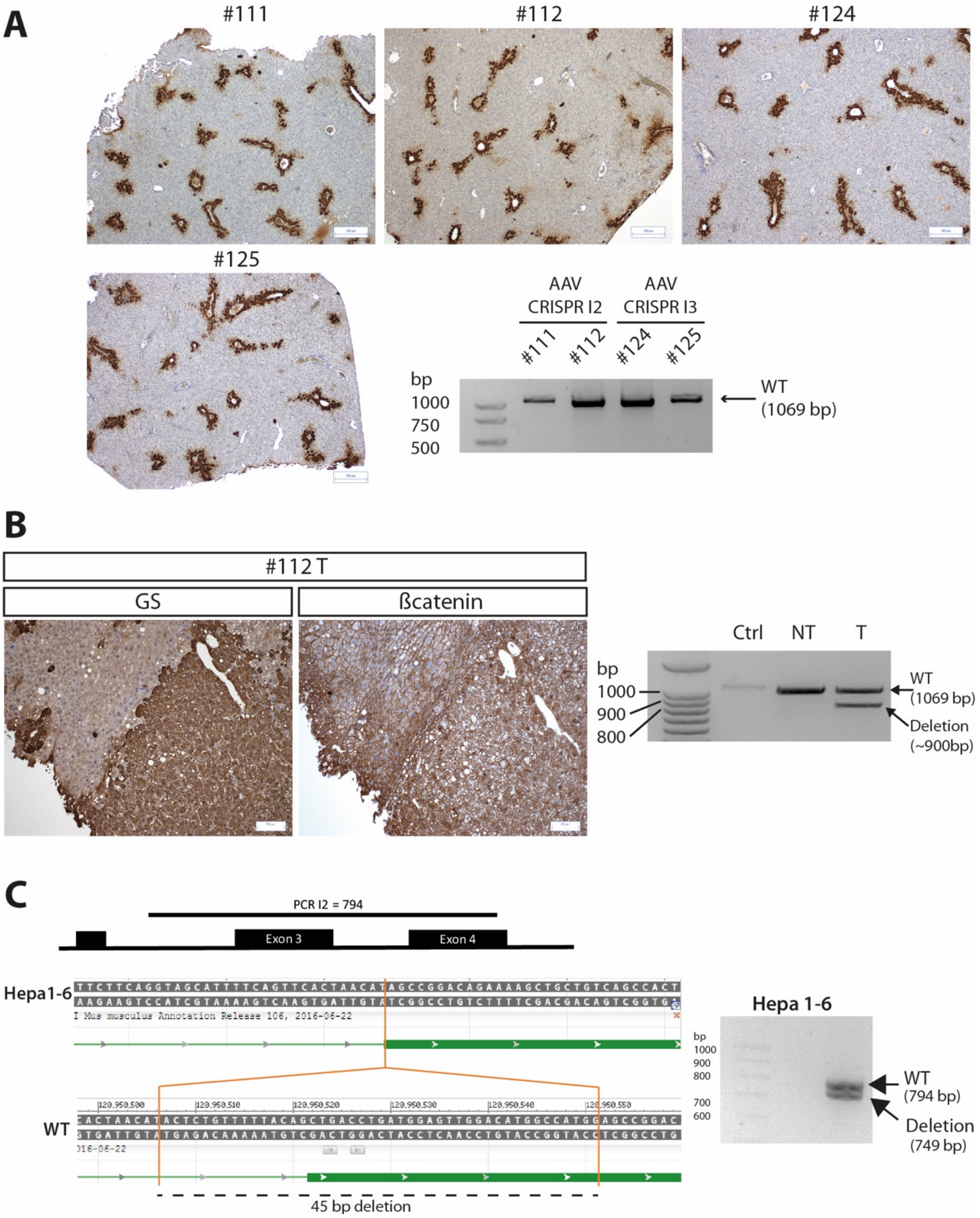


**Figure S5**: **AAV-CRISPR targeting of βcat-intron2 or βcat-intron3 in the liver. A.** GS IHC and ex3-PCR on liver biopsy two months after injection of $2.10^{10}$ vg of either sgI2 or sgI3. **B.** GS and β-catenin IHC reveal a β-catenin activated tumor that developed 8 months after injection of $2.10^{10}$ vg of sgI2 along with the corresponding ex3-PCR on non-tumoral (NT) and tumoral (T) tissue showing a >100bp deletion. **C.** Detection of a 45 bp deletion overlapping intron 2 and exon 3 in Hepa1-6 cell line via Sanger sequencing.

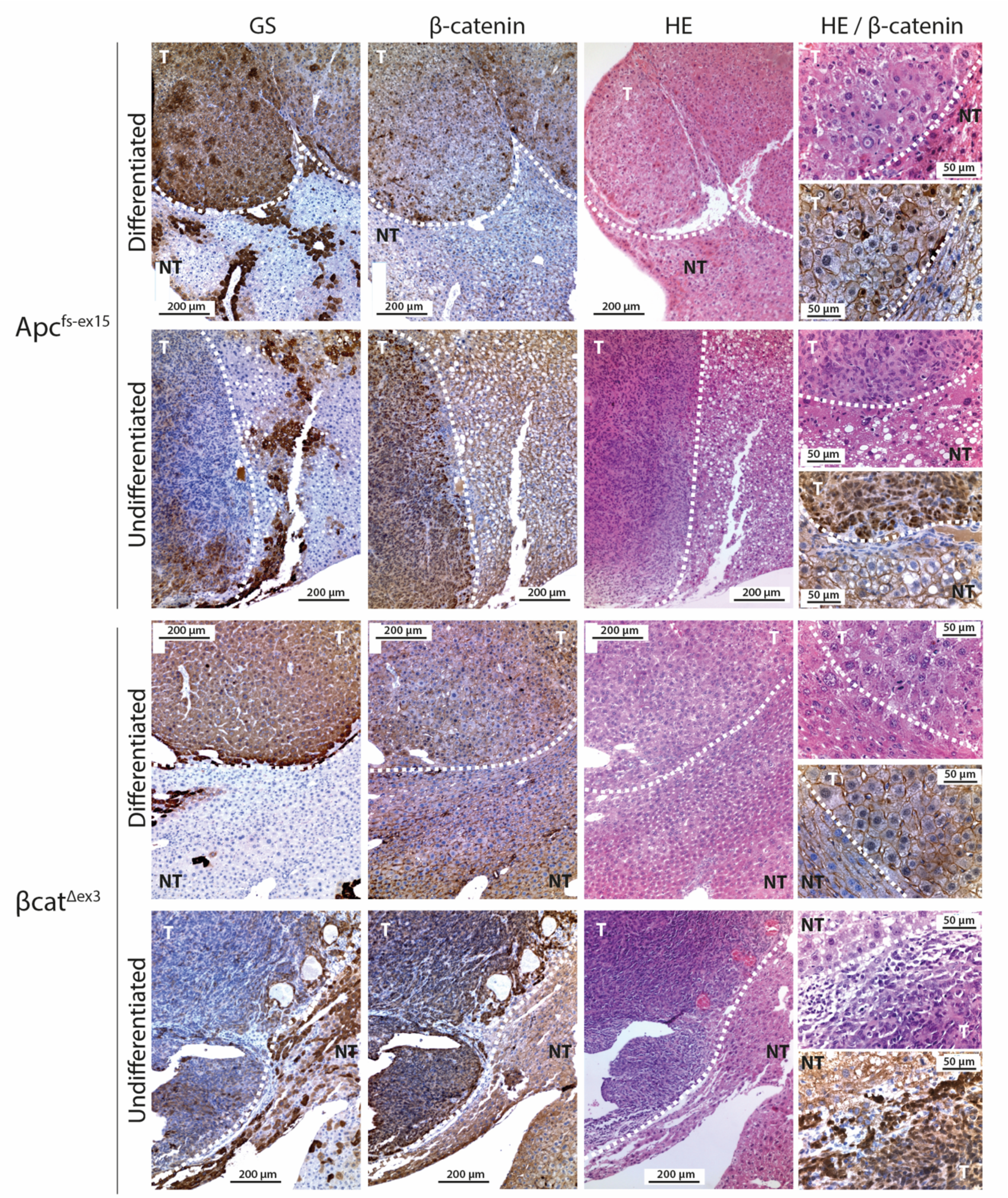


**Figure S6: *Apc*$^{fs\text{-}ex15}$ and βcat$^{\Delta ex3}$ tumors are phenotypically undistinguishable**. β-Catenin, GS and Hematoxyline-Eosine staining on representative differentiated and undifferentiated tumors from Crelox *Apc*$^{fs\text{-}ex15}$ and βcat$^{\Delta ex3}$ models.

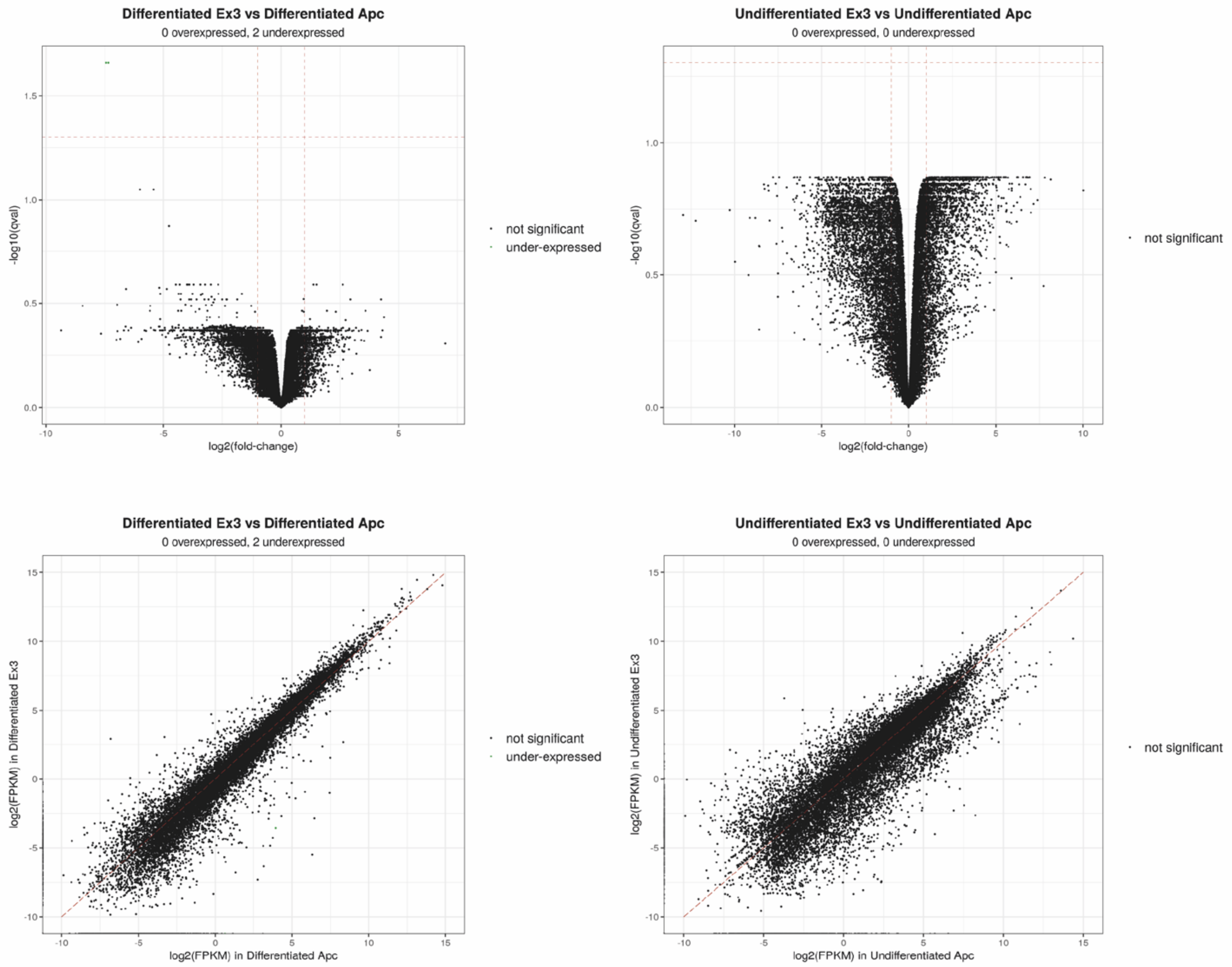


**Figure S7: volcano plot (Up) and scatter-plot (Down) of the differential analysis of Apc versus Ex3 tumors.**

| Tissue Type | n |
|---|---|
| Normal | 8 |
| HCC G1 | 8 |
| HCC G2 | 8 |
| HCC G3 | 8 |
| HCC G4 | 8 |
| HCC G5 | 8 |
| HCC G6 | 8 |
| HB Embrion. | 3 |
| HB Foetal | 3 |
| HB Mesench. | 3 |

| Genotype | n |
|---|---|
| APC KO | 6 |
| ΔEx3 | 5 |
| WT | 4 |

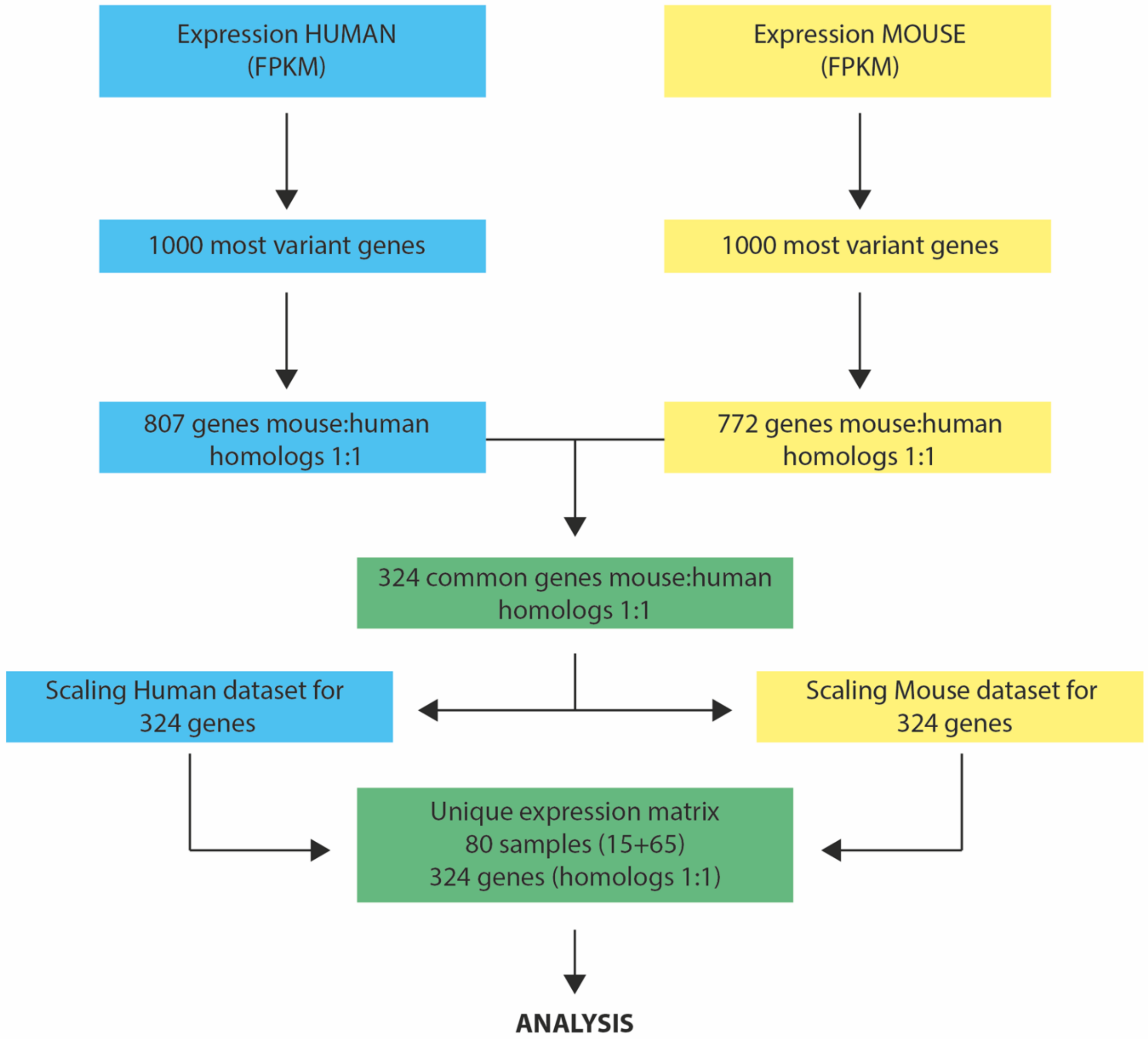


**Figure S8: Workflow for Integrated Analysis of Human (n=65)/Mouse (n=11) HCC/HB.**

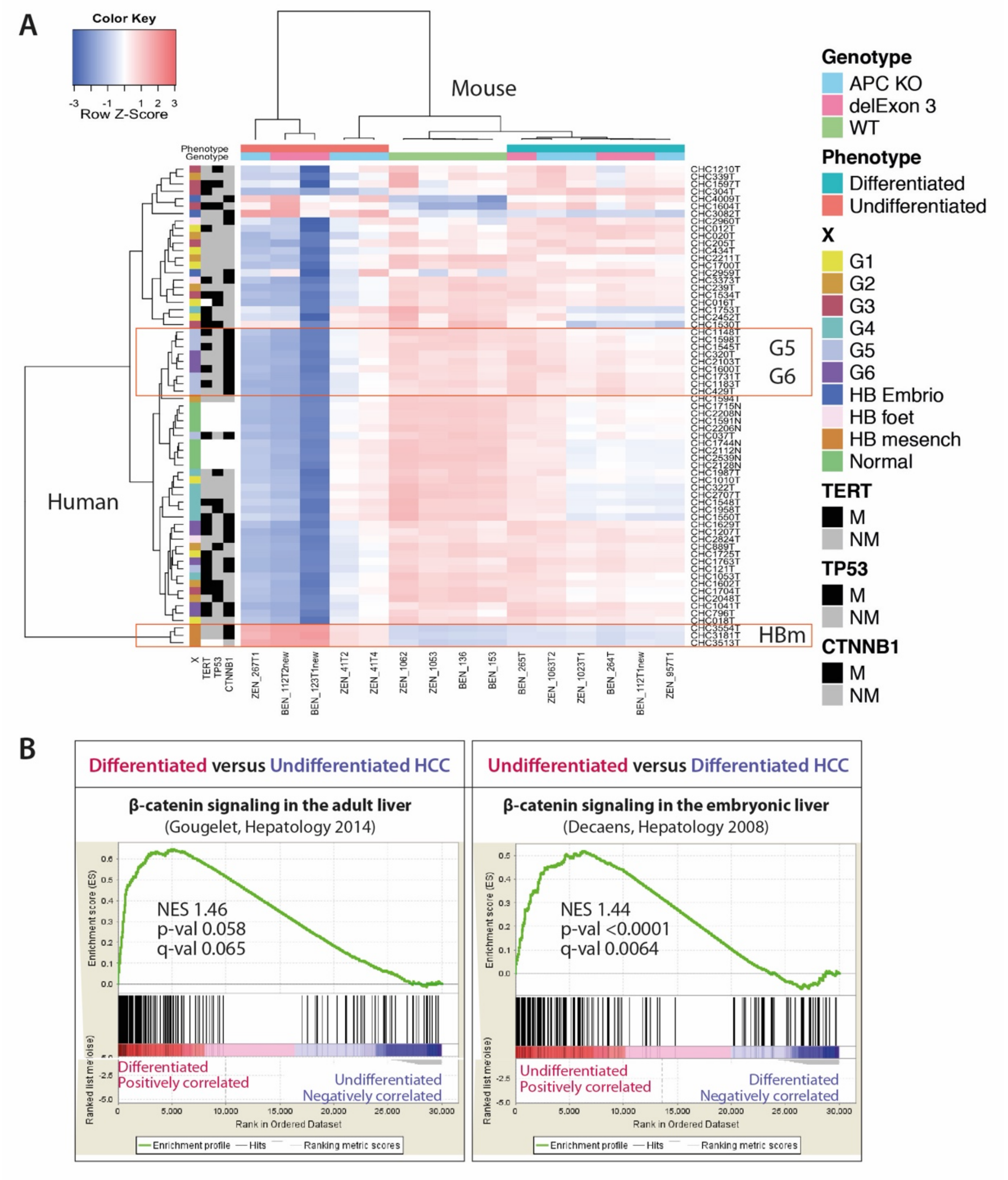


**Figure S9: Mouse undifferentiated tumors strongly correlate with human mesenchymal hepatoblastoma and a β-catenin signature in embryonic liver, while differentiated HCC correlate with G5-G6 classes of HCC and a β-catenin signature in adult liver. (A)** Correlation matrix; (B) GSEA using published gene datasets resulting from Apc loss and subsequent β-catenin activation in the adult or in the embryonic liver.

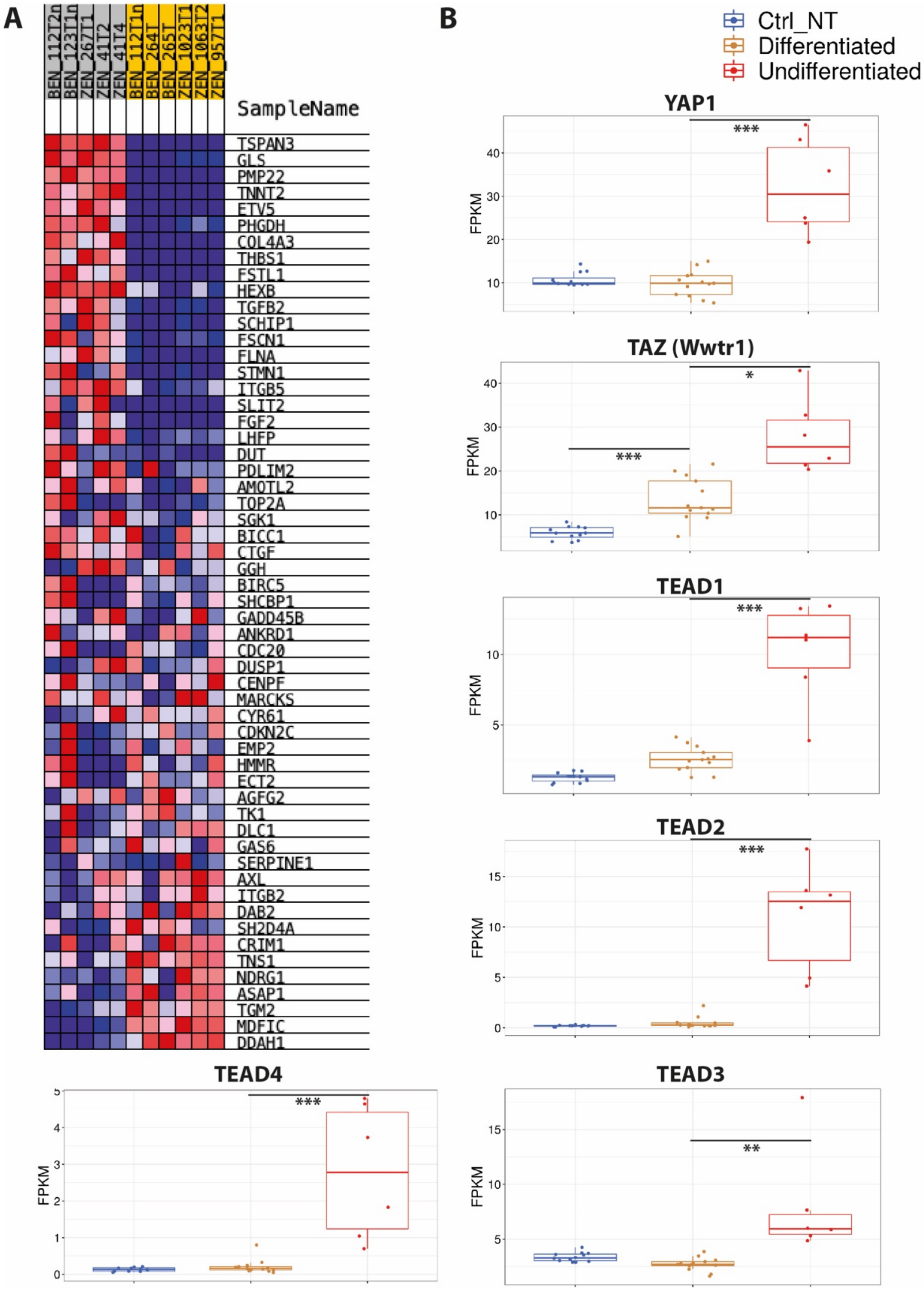


**Figure S10: YAP signature in undifferentiated mouse liver tumors.** RNASeq analyses: **A.** Overexpression of YAP signaling target genes (GSEA with "Cordenonsi YAP conserved signature"). **B.** Boxplots (Galileo Integragen) showing the overexpression of YAP signaling partners. *p<0.05, **p<0.01; ***p<0.001.

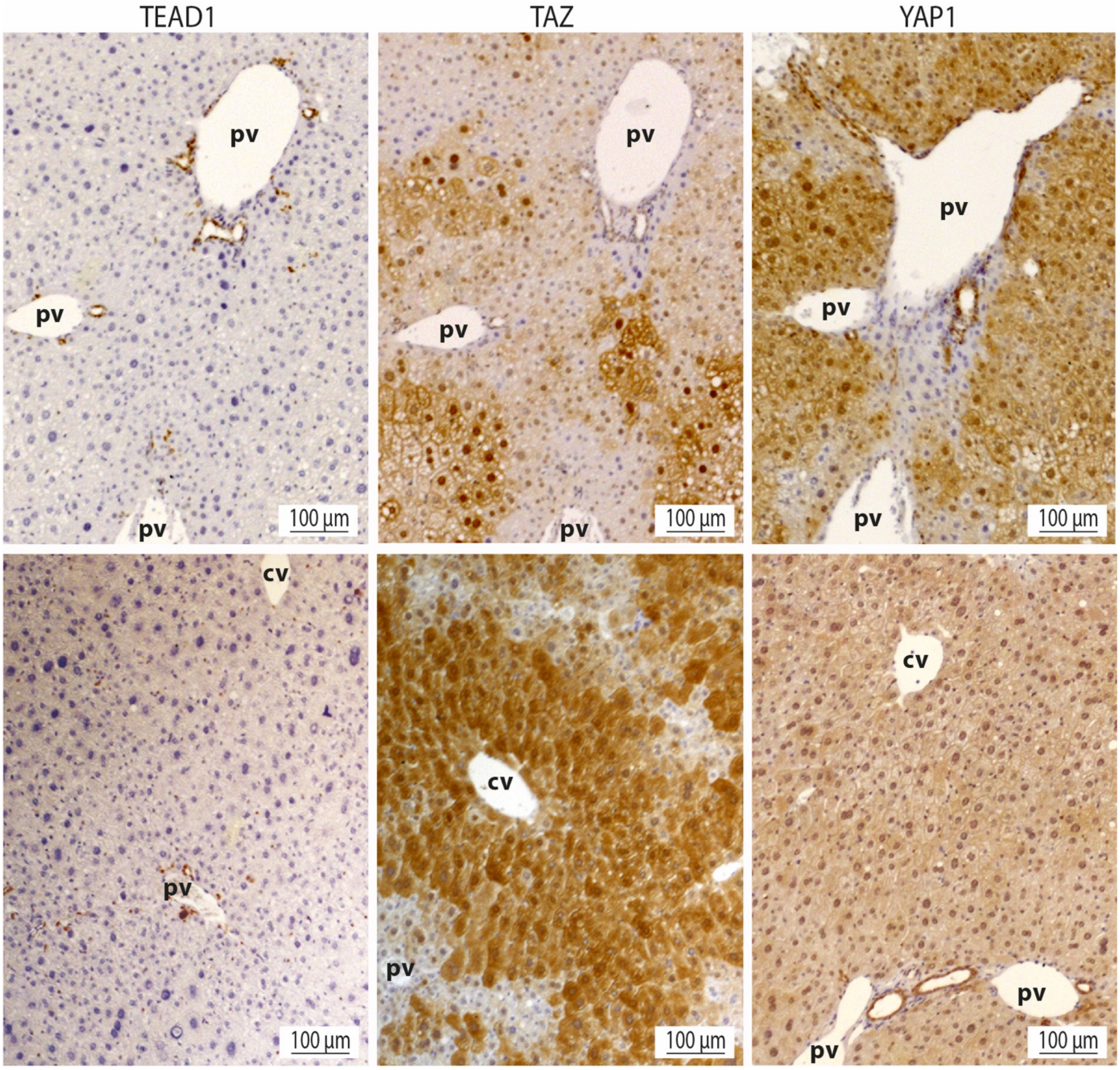


**Figure S11: Zonal expression of TEAD1, TAZ and YAP1 in normal livers.** Pv=portal vein, cv=centrolobular vein. TEAD1 expression is restricted to cholangiocytes into the bile ducts, near the portal vein. A mild TAZ expression is found in cholangiocytes, and a strong enrichment of both nuclear and cytosolic immunostaining is found in the pericentral hepatocytes. YAP1 expression is strong both in cholangiocytes and hepatocytes, and no apparent zonal expression can be seen.
